## Supplementary Materials for "Dating genomic variants and shared ancestry in population-scale sequencing data"

##### Contents

Supplementary Figures (S1–S9)
Supplementary Tables (S1–S2)
Supplementary Text

### Supplementary Figures

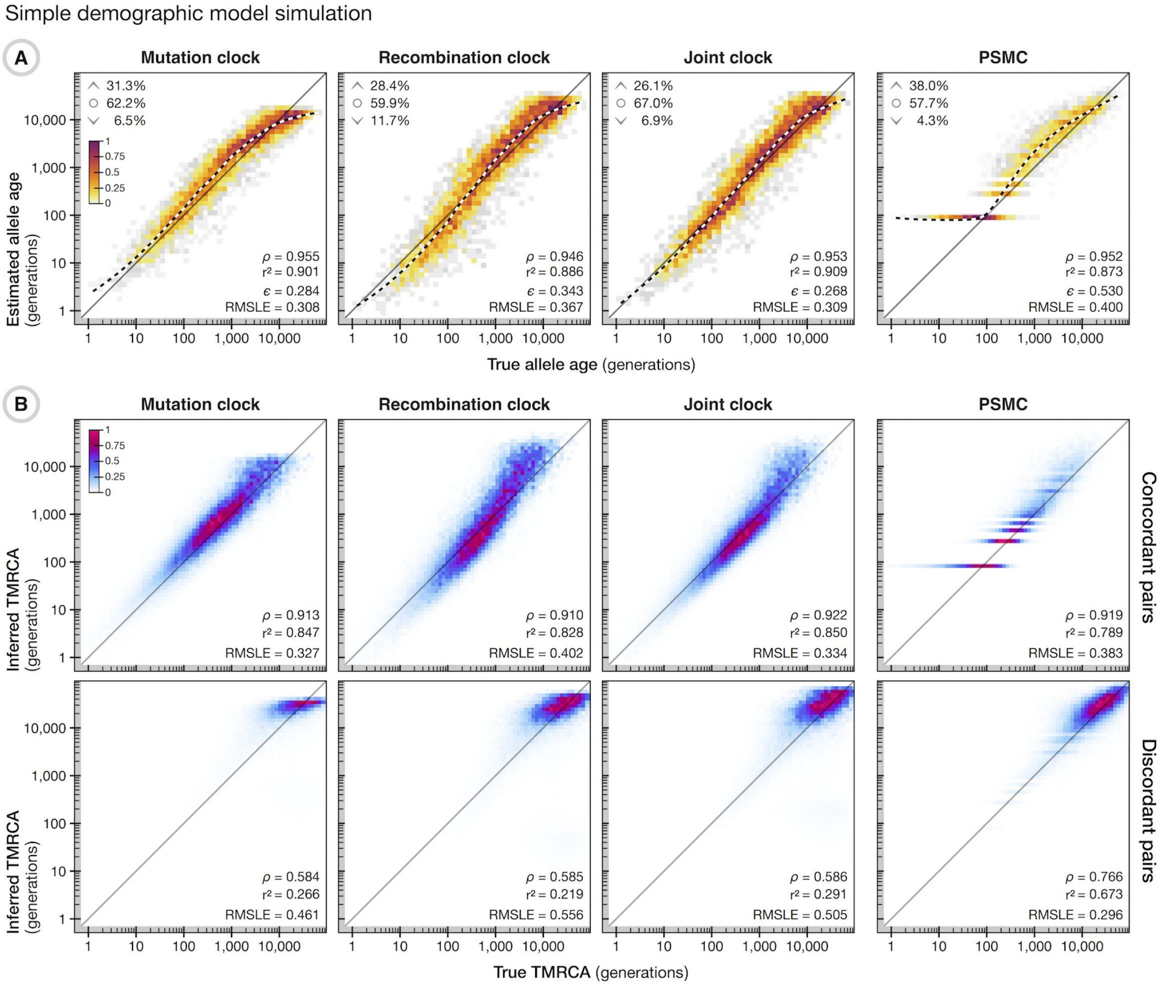

**Figure S1. Simple demographic model simulation.** Allele age estimation and TMRCA inference in a simulation of a 100 Mb region with sample size $N$=1,000, effective population size $N_e$=10,000, constant mutation rate ($\mu = \times 10^{-8}$ per site per generation), and constant recombination rate ($r = 1 \times 10^{-8}$ per site per generation). (**A**) Relationship between the true allele age (geometric mean of lower and upper age of the branch on which a mutation occurred; *x-axis*) and estimated allele age (*y-axis*), estimated using the mutation clock, recombination clock, and joint clock models (*left*) and PSMC (*right*) for the same set of 5,000 variants, randomly sampled at allele count $1 < x < N$. Colors indicate the density scaled by the maximum per panel. Upper inserts indicate the fraction of sites where the point estimate (mode of the composite posterior distribution) of allele age lies above the upper age of the branch on which the mutation occurred ($\wedge$), below the lower age ($\vee$), or within the range of the branch ($\circ$). Lower inserts indicate the Spearman rank correlation statistic, $\rho$, the square of the Pearson correlation coefficient (on log-scale), $r^2$, the interval-adjusted error metric, $\epsilon$, and the root mean squared $\log_{10}$ error, RMSLE. Also shown is a LOESS fit (2nd degree polynomials, neighborhood proportion $\alpha = 0.25$; *dashed line*). (**B**) Relationship between the true TMRCA for a haplotype pair at a given site and the corresponding inferred TMRCA (mean of posterior distribution), shown separately for concordant and discordant pairs. The same sets of haplotype pairs were analyzed under each clock model in GEVA (*left*) and PSMC (*right*). Colors indicate the density scaled by the maximum per panel. Lower inserts indicate the Spearman rank correlation statistic, $\rho$, the square of the Pearson correlation coefficient (on log-scale), $r^2$, and the root mean squared $\log_{10}$ error, RMSLE.

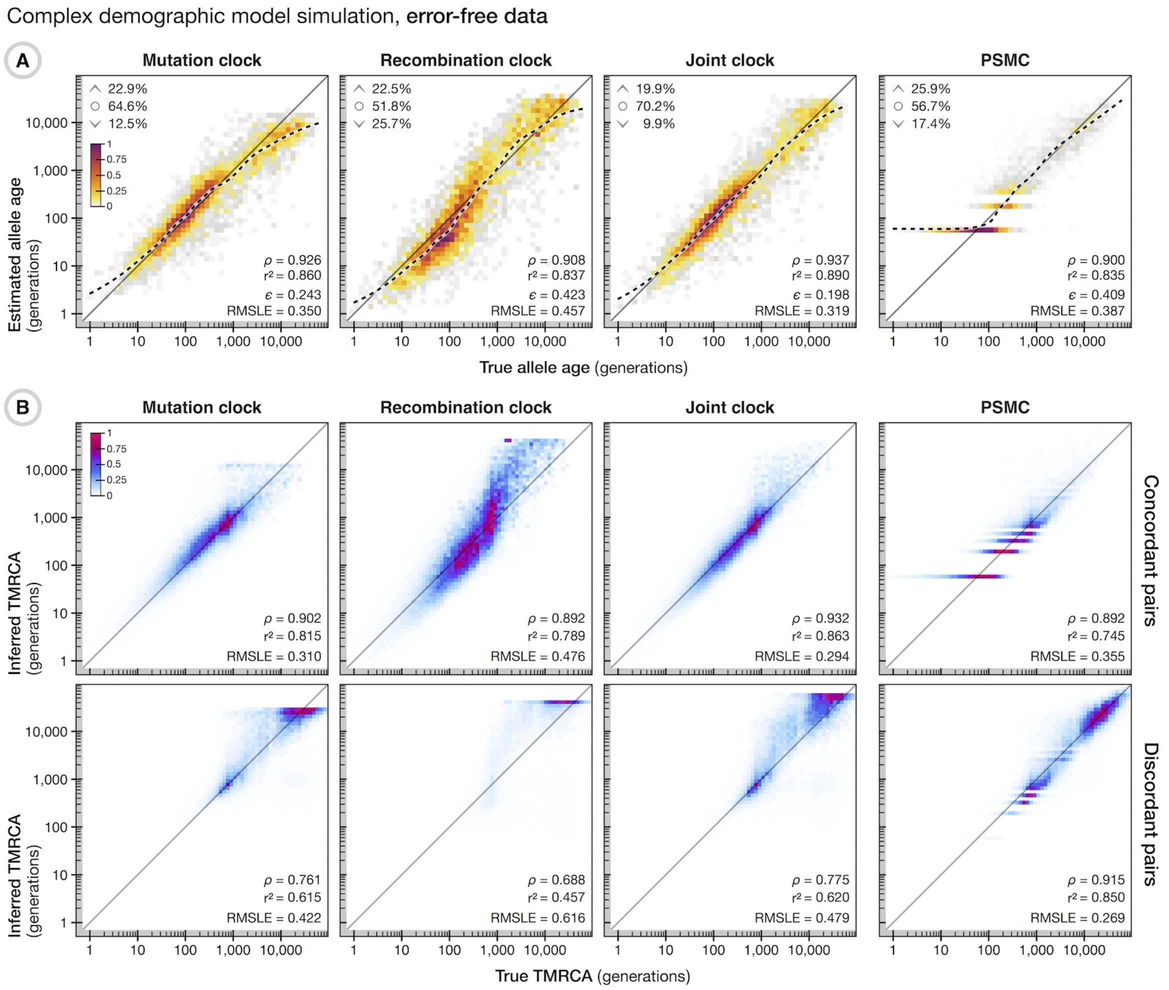

**Figure S2. Complex demographic model simulation without error.** Allele age estimation and TMRCA inference in a simulation that recapitulates the human expansion out of Africa [1], with $N$=5,000, $N_e$=7,300, constant mutation rate $\mu = 2.35 \times 10^{-8}$, and variable recombination rates from HapMap (Phase 2, GRCh37) [2] for Chromosome 20 (63 Mb). Allele age was estimated for 5,000 variants sampled uniformly from the intersection of sites available at allele count $1 < x < N$ in data without error and after data were modified with error; see Figures S3 and S4. (**A**) See description in Figure S1A. (**B**) See description in Figure S1B.

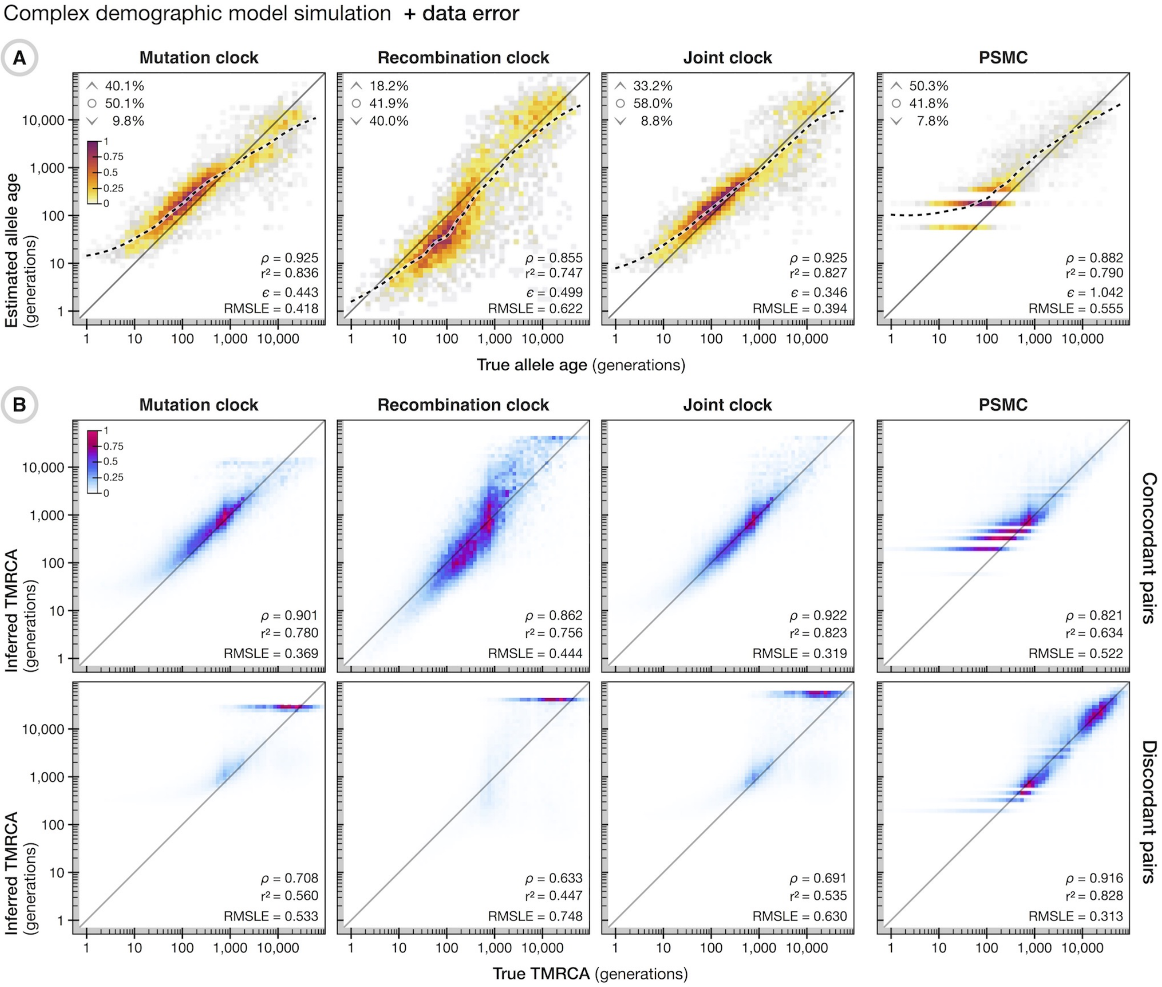

**Figure S3. Complex demographic model simulation with error.** Allele age estimation and TMRCA inference from simulated data in which haplotype data were modified with realistic error rates; calibrated from empirical estimates of genotype errors in 1000 Genomes Project data [3], by comparison to corresponding genotype data from the Illumina Platinum Genomes Project [4]. Allele age was estimated for the same set of 5,000 variants as analyzed in Figures S2–S4. (**A**) See description in Figure S1A. (**B**) See description in Figure S1B.

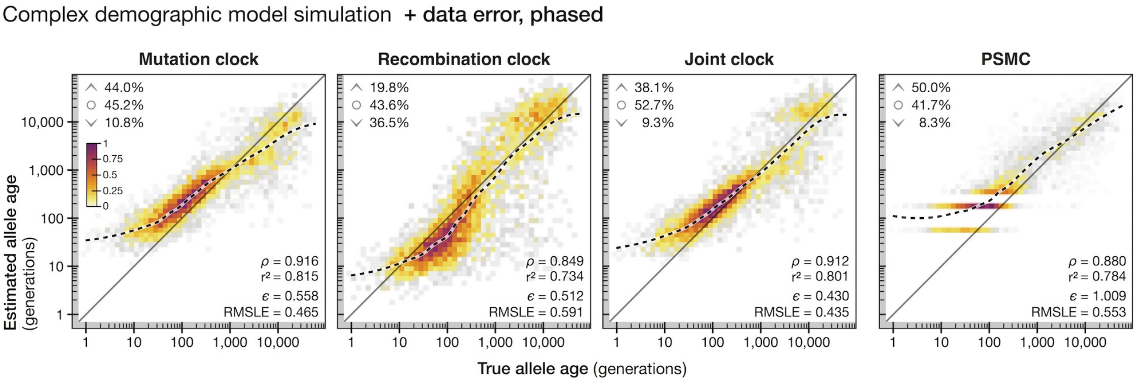

**Figure S4. Complex demographic model simulation with error and after phasing.** Allele age estimation and TMRCA inference from simulated data in which haplotype data were modified with realistic error rates; calibrated from empirical estimates of genotype errors in 1000 Genomes Project data [3], by comparison to data from the Illumina Platinum Genomes Project [4]. Haplotype data were additionally phased using `SHAPEIT2` [5] after the introduction of data error. Allele age was estimated for the same set of 5,000 variants as analyzed in Figures S2 and S3. Description of plots as in Figure S1A. Note that the relationship between true and inferred TMRCA per haplotype pair (as in Figures S1–S3) cannot be ascertained conclusively after phasing of haplotype data.

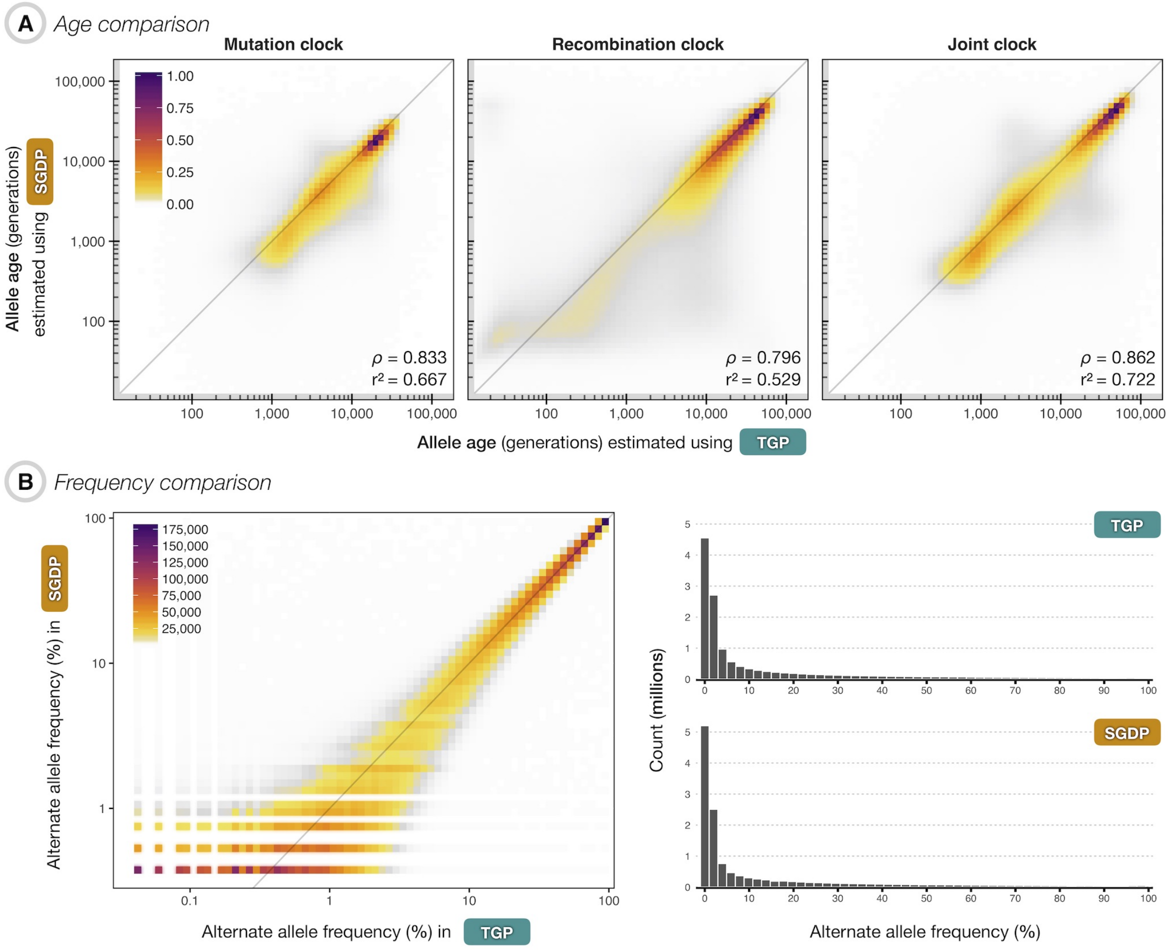

**Figure S5. Correlation between allele age estimated separately in TGP and SGDP data.** (**A**) The relationship between allele age using data from TGP (*x-axis*) and SGDP (*y-axis*), estimated from the mutation clock (*left*), recombination clock (*center*), and the joint clock model (*right*), for 13.7 million variants dated in both data sources. Colors indicate the density scaled by the maximum per panel. Lower inserts indicate the Spearman rank correlation statistic, $\rho$, and the square of the Pearson correlation coefficient (calculated on log-scaled allele ages), $r^2$. (**B**) Differences in allele frequency of the variants compared (*left*); the histograms (*right*) show the frequencies as observed in TGP (*top*) and SGDP (*bottom*) for corresponding sets of variants.

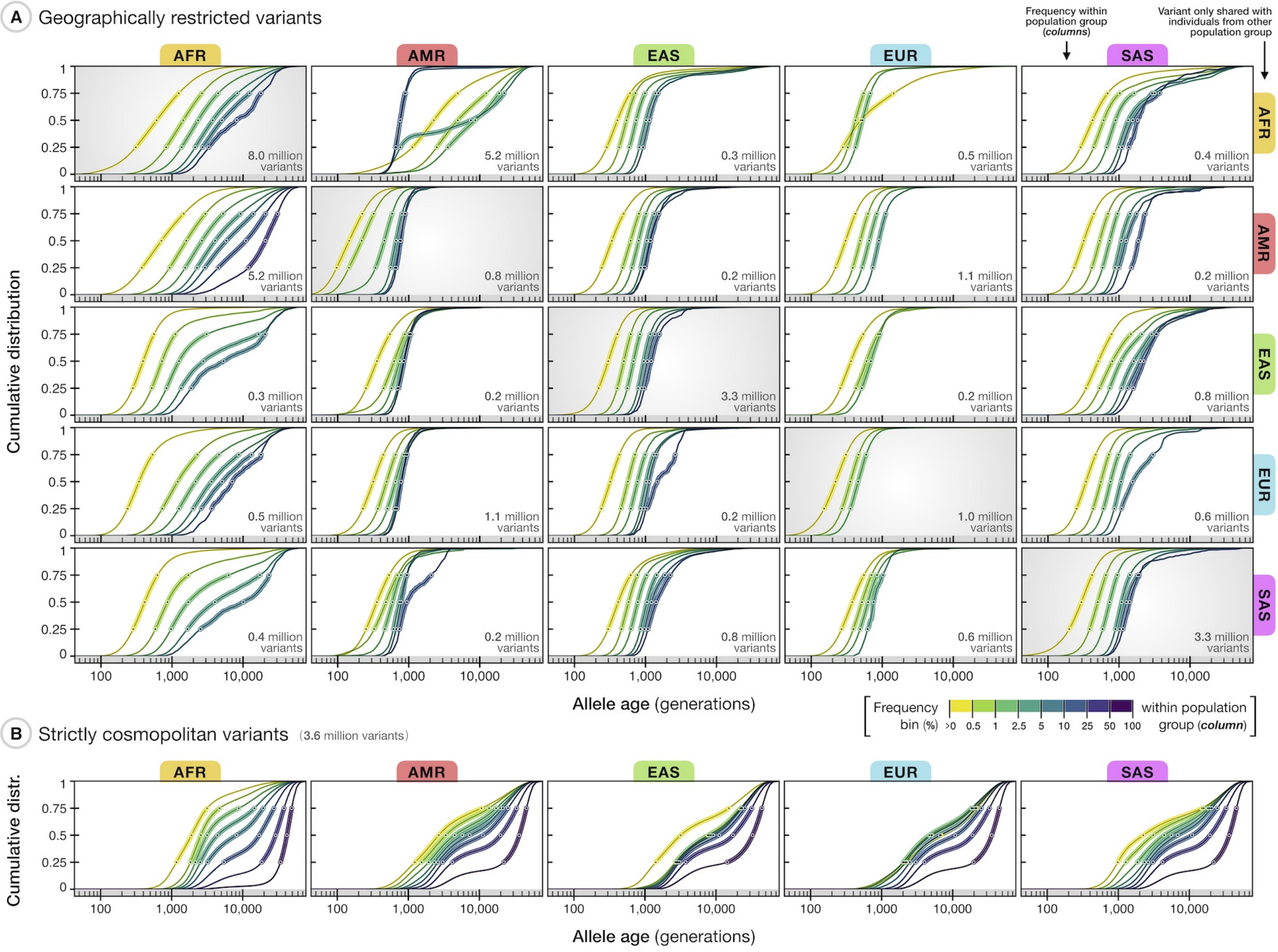

**Figure S6. Allele age and frequency for variants shared between different human populations.** The relationship between allele age and frequency for variants dated in from the 1000 Genomes Project (TGP). Allele age was estimated under the joint clock model using TGP data (whole sample). Of the 43.2 million variants dated in TGP (Chromosomes 1-22), we excluded those with low estimation quality and inconsistent ancestral allele information, which retained 34.4 million variants. Allele frequencies were calculated within subsamples of African (AFR), American (AMR), East Asian (EAS), European (EUR), and South Asian (SAS) ancestry groups, as defined in TGP sample data. Lines in each panel show the cumulative age distribution of variants within a given frequency bin (see legend), with frequencies as observed within the population group indicated at the top (*columns*); circles indicate median and inter-quartile range (25th, 50th, and 75th percentiles). (**A**) The subset of variants observed in only two population groups, referring to sites that have non-zero frequencies in either of the two populations considered and zero frequency in all other groups (*white* background color), and geographically restricted variants that are isolated within only a single population group, referring to sites that have non-zero frequencies in the population considered and zero frequency in all other groups (diagonal panels; *grey* background color). Panels that are diagonal opposites show results for the same set of variants, but with frequencies as observed in the population considered (by column). The number of variants retained is shown in each panel (*bottom right*). (**B**) The age distribution of strictly cosmopolitan variants (non-zero frequencies in every group) by frequency as seen within a population group. The distributions shown in each panel were obtained on same set of 3,634,716 variants. A more detailed summary for geographically restricted variants (those that are isolated within a given group; A, diagonal panels) and cosmopolitan variants (B) is given in Table S2.

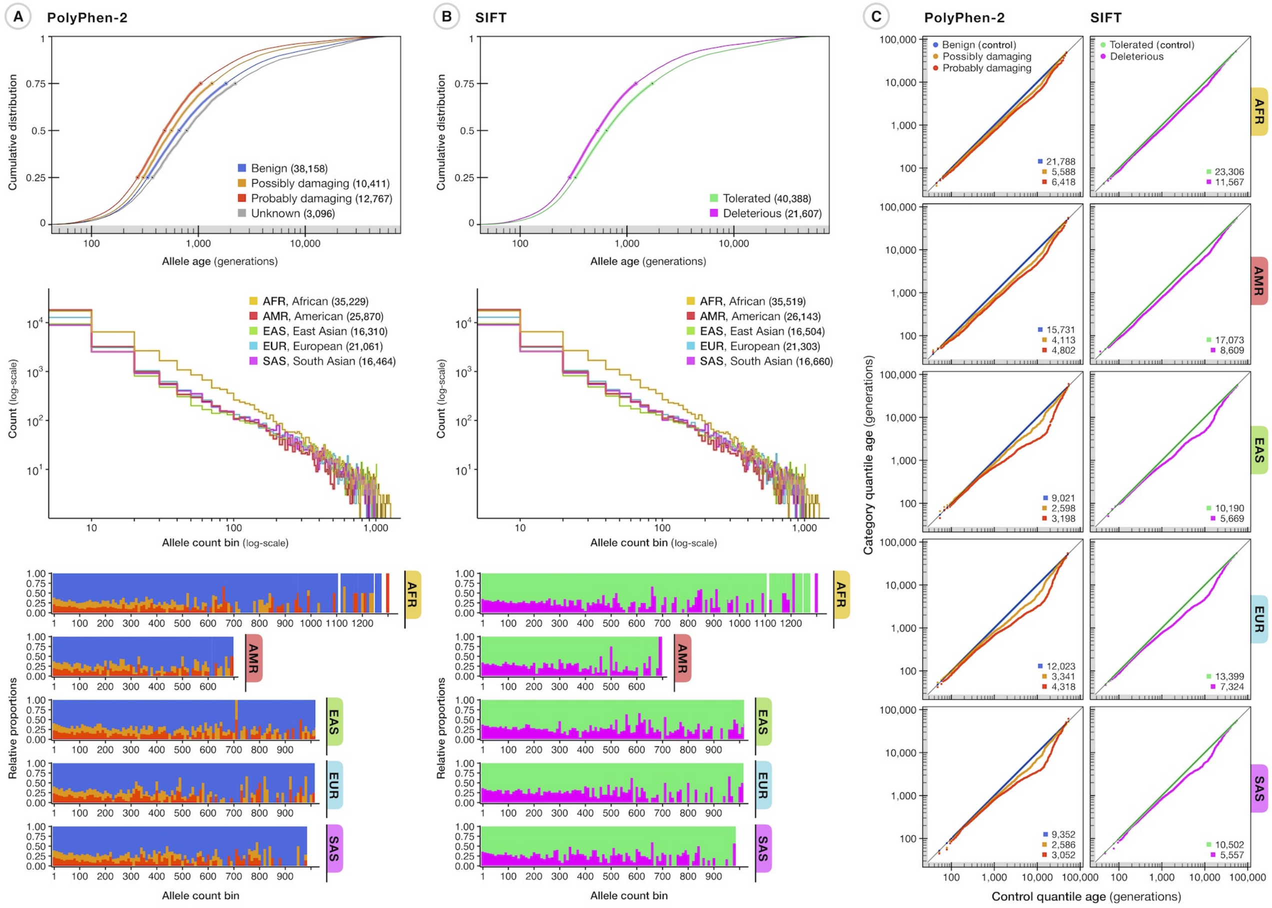

**Figure S7. Allele age of potentially pathogenic variants.** Allele age was estimated (joint clock) using sample data from the 1000 Genomes Project (TGP) for variants annotated by the Ensembl Variant Effect Predictor (VEP) [6]. We excluded variants with low age estimation quality or inconsistent ancestral allelic states (determined through multi-species alignments; information available through Ensembl), which retained 64,432 (of 70,220) variants annotated by PolyPhen-2 [7] and 61,995 (of 67,539) variants annotated by SIFT [8]; of those, 61,615 (of 67,123) variants have been annotated by both methods. (**A**) The relationship between allele age and variant effects predicted by PolyPhen-2 (*top*), with effect categories given as *benign*, *possibly damaging*, *probably damaging*, and *unknown*; variant numbers per category are indicated in the legend. Each line shows the cumulative age distribution by effect category; circles indicate median and inter-quartile range. The frequency distribution of all variants considered is shown for the five major population groups defined in the TGP sample (*middle*), given as the number of variants within allele count bins (evenly spaced on linear scale but shown on log scale), with allele count as per population group in TGP. The number of variants at non-zero frequencies in a population group is indicated in the legend. The relative proportion of variants across effect categories per allele count bin for the five major population groups in TGP (*bottom*). (**B**) As for part (A), plots show the relationships between allele age and population frequency for variants predicted by SIFT, with effect categories given as *tolerated* and *deleterious*. (**C**) QQ-plots showing differences in allele age distributions for variants annotated by PolyPhen-2 (*left*) and SIFT (*right*), compared to a control set of variants (those annotated as *benign* by PolyPhen-2 or *tolerated* by SIFT), matched for allele frequency within a given population group. Matching was done by retaining only those variants observed at non-zero frequency within a population group, and if variants of every effect category were represented at identical allele counts. The insert in each panel (*bottom right*) shows the number of variants retained per effect category.

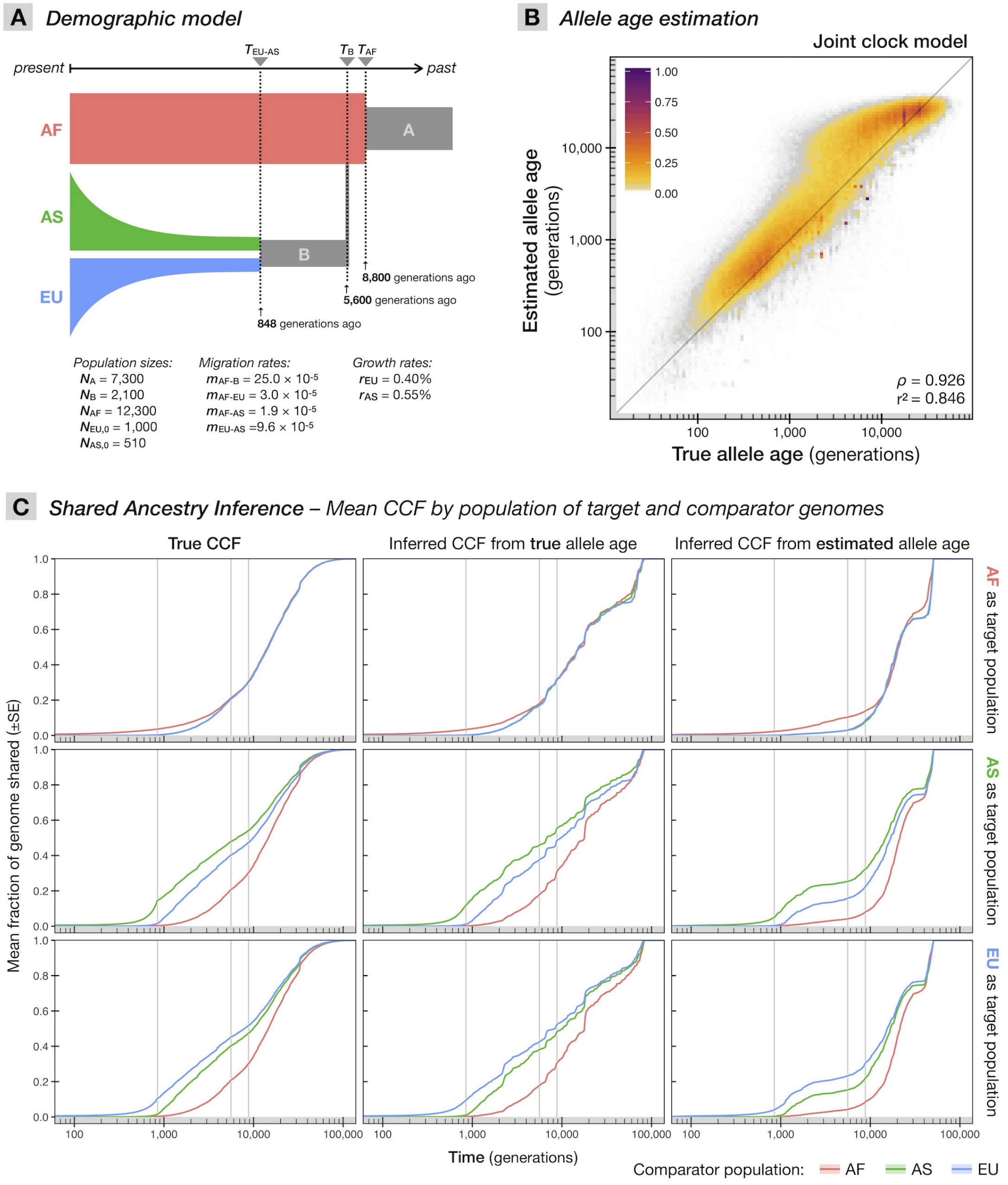

**Figure S8. Shared ancestry inference in simulated data.** (**A**) The demographic model used in coalescent simulations, which recapitulates the human expansion out of Africa [1], for three major populations; African (AF), Asian (AS), and European (EU). The times of three events are indicated (*top*); $T_{AF}$ marks the time when AF emerged from ancestral population A, $T_B$ the time when ancestral population B emerged from AF, and $T_{EU-AS}$ the split of EU and AS from B. The time of each event assumes a generation time of 25 years per generation. The simulation was conducted with parameters as stated in the figure (*bottom*); that is, the size of each population, where $N_{AS,0}$ and $N_{EU,0}$ refer to the initial size of AS and EU, respectively, migration rates, and growth rates for AS and EU. We used this model to simulate a sample of $N = 600$ haplotypes (200 haplotypes from each of the three populations). (**B**) GEVA estimates of 412,344 variants (all variants at allele count $1 < x < N$), comparing true age (geometric mean of lower and upper age of the branch on which a mutation occurred; *x-axis*) and estimated age (joint clock model, $\max_{\mathcal{C}} = 100$, $\max_{\mathcal{D}} = 100$; *y-axis*). Colors indicate the maximized density; the insert shows the Spearman rank correlation statistic, $\rho$, and the square of the Pearson correlation coefficient (calculated on log-scaled age), $r^2$. (**C**) Shared ancestry inference, comparing coalescent profiles obtained from simulation records (true CCF; *left*), inferred from true allele age (*center*), and inferred from estimated allele age (*right*). We computed the CCFs for each of the 600 simulated haplotypes as target genome in turn against the whole sample as comparators; recorded over a fixed grid of 500 time points between 1 and 500,000 generations ago equally spaced on log-scale. Each line shows the mean and standard error (±SE) of CCFs between each combination of target and comparator population; for target genomes from AF (*top*), AS (*middle*), and EU (*bottom*). Vertical lines indicate the times of the three demographic events; $T_{EU-AS}$ (*left*), $T_B$ (*center*), and $T_{AF}$ (*right*). These results are discussed in further detail in the Supplementary Text (Section S6.2).

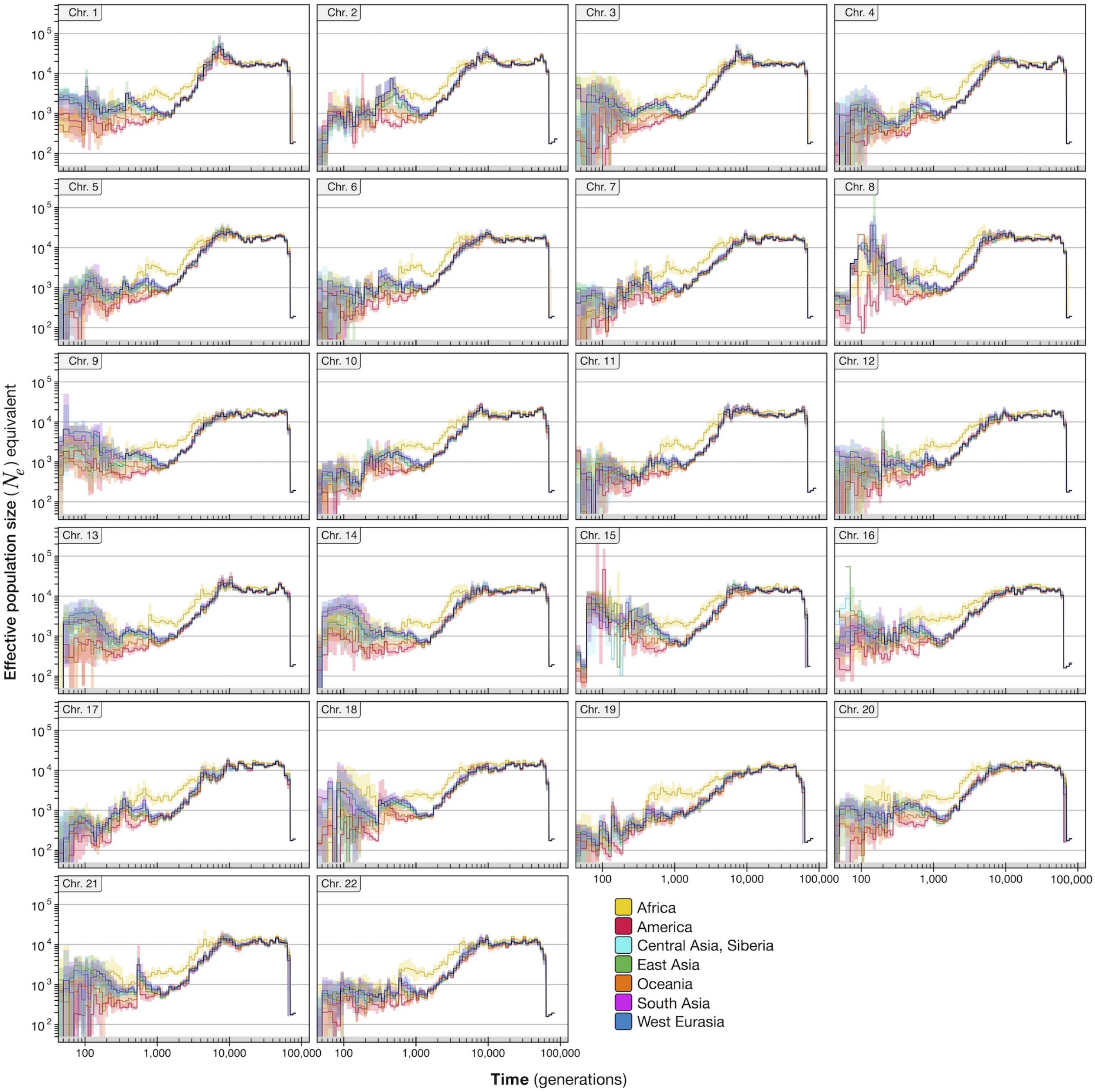

**Figure S9. Effective population size ($N_e$ equivalent) over time for Chromosomes 1-22 in SGDP.** The cumulative coalescent function (CCF) was inferred for each haploid target genome with all other comparator genomes in the SGDP sample, based on a total of 11.7 million variants dated (joint clock) on Chromosomes 1-22, retained after excluding variants with low age estimation quality and inconsistent ancestral allele information. Coalescent intensity was computed per target genome and scaled by the maximum over the sample at a given time interval (epoch; evenly distributed on log-scale). Each line shows the median and inter-quartile range of $N_e$ equivalents inferred for individuals in the different ancestry groups (continental regions; see legend).

### Supplementary Tables

**Table S1. Summary of variants per chromosome in the Atlas of Variant Age.** The table shows the total number ($N_{\text{all}}$) of variants available in the Atlas of Variant Age on Chromosomes 1-22, as well as the number of variants dated using data from the 1000 Genomes Project (TGP) alone ($N_{\text{TGP}}$) and the Simons Genome Diversity Project (SGDP) alone ($N_{\text{SGDP}}$). Additionally, variants present in both data sets were dated using the independently inferred pairwise TMRCA results from TGP and SGDP to obtain a *combined* age estimate ($N_{\text{Combined}}$). The number of haplotype pairs at which shared haplotype segments and TMRCA were inferred is shown for the two data sources; numbers are shown as the *sum* of *concordant* and *discordant* pairs analyzed per chromosome. See Section S7 for details about the analysis of TGP and SGDP sample data. Full result data sets for each variant, including the results of each pairwise analysis and age estimates obtained under each clock model (mutation, recombination, and joint clock; see Section S1.3), are publicly available online at `https://human.genome.dating`.

| Chr. | $N_{\text{all}}$ | $N_{\text{TGP}}$ | $N_{\text{SGDP}}$ | $N_{\text{Combined}}$ | Haplotype pairs analyzed in **TGP** | | | Haplotype pairs analyzed in **SGDP** | | |
|---|---|---|---|---|---|---|---|---|---|---|
| | | | | | Sum = | Concordant + | Discordant | Sum = | Concordant + | Discordant |
| 1 | 3,603,845 | 3,434,810 | 1,242,878 | 1,073,843 | 2,361,753,523 = | 670,234,719 + | 1,691,518,804 | 178,223,566 = | 57,340,106 + | 120,883,460 |
| 2 | 3,909,601 | 3,727,112 | 1,354,976 | 1,172,487 | 2,552,046,545 = | 717,105,655 + | 1,834,940,890 | 193,049,416 = | 61,961,663 + | 131,087,753 |
| 3 | 3,246,604 | 3,078,989 | 1,184,447 | 1,016,832 | 2,121,176,886 = | 604,531,045 + | 1,516,645,841 | 170,540,962 = | 55,202,026 + | 115,338,936 |
| 4 | 3,207,690 | 3,055,978 | 1,156,613 | 1,004,901 | 2,122,460,565 = | 616,887,880 + | 1,505,572,685 | 167,838,575 = | 55,197,442 + | 112,641,133 |
| 5 | 2,936,062 | 2,777,306 | 1,054,634 | 895,878 | 1,909,734,447 = | 541,892,101 + | 1,367,842,346 | 151,025,979 = | 48,366,615 + | 102,659,364 |
| 6 | 2,841,180 | 2,703,481 | 1,023,730 | 886,031 | 1,883,434,856 = | 551,467,803 + | 1,331,967,053 | 148,183,411 = | 48,575,536 + | 99,607,875 |
| 7 | 2,651,520 | 2,530,710 | 895,609 | 774,799 | 1,748,759,380 = | 502,479,763 + | 1,246,279,617 | 129,367,159 = | 42,116,601 + | 87,250,558 |
| 8 | 2,574,597 | 2,446,808 | 918,980 | 791,191 | 1,681,198,356 = | 475,982,722 + | 1,205,215,634 | 131,959,407 = | 42,552,249 + | 89,407,158 |
| 9 | 1,992,571 | 1,898,132 | 695,180 | 600,741 | 1,306,395,549 = | 371,461,422 + | 934,934,127 | 100,294,995 = | 32,686,939 + | 67,608,056 |
| 10 | 2,246,011 | 2,145,420 | 786,737 | 686,146 | 1,484,038,224 = | 426,770,806 + | 1,057,267,418 | 114,586,646 = | 37,957,269 + | 76,629,377 |
| 11 | 2,241,144 | 2,144,729 | 758,739 | 662,324 | 1,477,906,889 = | 421,948,241 + | 1,055,958,648 | 109,601,162 = | 35,707,665 + | 73,893,497 |
| 12 | 2,152,949 | 2,051,957 | 747,059 | 646,067 | 1,418,690,735 = | 407,634,945 + | 1,011,055,790 | 108,059,559 = | 35,361,194 + | 72,698,365 |
| 13 | 1,595,057 | 1,517,081 | 591,225 | 513,249 | 1,052,112,658 = | 304,833,376 + | 747,279,282 | 85,982,601 = | 28,409,968 + | 57,572,633 |
| 14 | 1,477,988 | 1,406,101 | 523,596 | 451,709 | 970,068,468 = | 277,776,895 + | 692,291,573 | 75,600,107 = | 24,647,165 + | 50,952,942 |
| 15 | 1,351,709 | 1,287,260 | 466,764 | 402,315 | 885,168,969 = | 251,621,540 + | 633,547,429 | 67,128,688 = | 21,737,994 + | 45,390,694 |
| 16 | 1,501,059 | 1,431,898 | 497,060 | 427,899 | 976,581,337 = | 271,471,963 + | 705,109,374 | 71,332,335 = | 22,986,825 + | 48,345,510 |
| 17 | 1,290,584 | 1,230,848 | 416,535 | 356,799 | 840,095,235 = | 234,006,858 + | 606,088,377 | 59,898,160 = | 19,313,408 + | 40,584,752 |
| 18 | 1,275,859 | 1,212,338 | 477,771 | 414,250 | 838,516,668 = | 241,202,386 + | 597,314,282 | 69,122,568 = | 22,576,114 + | 46,546,454 |
| 19 | 1,028,005 | 993,091 | 252,473 | 217,559 | 685,952,950 = | 196,580,072 + | 489,372,878 | 36,536,015 = | 11,914,815 + | 24,621,200 |
| 20 | 1,016,303 | 964,721 | 363,883 | 312,301 | 663,506,829 = | 187,934,142 + | 475,572,687 | 52,243,054 = | 16,822,189 + | 35,420,865 |
| 21 | 628,110 | 596,192 | 226,055 | 194,137 | 412,235,218 = | 118,511,806 + | 293,723,412 | 32,855,224 = | 10,805,152 + | 22,050,072 |
| 22 | 625,257 | 597,558 | 199,880 | 172,181 | 413,241,926 = | 119,074,286 + | 294,167,640 | 28,956,345 = | 9,502,483 + | 19,453,862 |
| **Total** | **45,393,705** | 43,232,520 | 15,834,824 | 13,673,639 | **29,805,076,213** = | 8,511,410,426 + | 21,293,665,787 | **2,282,385,934** = | 741,741,418 + | 1,540,644,516 |

**Table S2. Age and frequency of variants within population groups in 1000 Genomes Project (TGP) data.** We estimated allele age for variants identified in TGP to characterize the age distribution of genetic variation across the human genome. Allele age was estimated under the joint clock model. Of the 43,232,520 variants dated in TGP (Chromosomes 1-22), we retained only those at quality score $QS > 0.5$ (see Section S1.5.1) and at which the ancestral allele is known and mapped to the reference allele (see Section S7.1), which retained 34,388,511 variants. The table shows the number of variants ($N$) and the median of allele age estimates ($Q_{50}$), as well as the 25th ($Q_{25}$) and 75th ($Q_{75}$) percentiles, per continental population group and stratified by allele frequency within that group. This is shown for (**A**) variants at non-zero frequencies within a given ancestry group, (**B**) geographically restricted variants that segregate only within a given group, and (**C**) strictly cosmopolitan variants that are shared among individuals from every continental group. Populations are abbreviated as follows; AFR: African; AMR: American; EAS: East Asian; EUR: European; SAS: South Asian.

| Population group | Frequency range (%) | (**A**) Frequency within population | | | | (**B**) Geographically restricted variants | | | | (**C**) Strictly cosmopolitan variants | | | |
|---|---|---|---|---|---|---|---|---|---|---|---|---|---|
| | | $N$ | $Q_{25}$ | $Q_{50}$ | $Q_{75}$ | $N$ | $Q_{25}$ | $Q_{50}$ | $Q_{75}$ | $N$ | $Q_{25}$ | $Q_{50}$ | $Q_{75}$ |
| **AFR** | $(0, 0.5]$ | 8,933,432 | 351 | 670 | 1,287 | 5,140,255 | 309 | 607 | 1,244 | 476,121 | 1,169 | 1,902 | 3,155 |
| | $(0.5, 1.0]$ | 2,699,024 | 886 | 1,558 | 2,857 | 1,536,213 | 826 | 1,439 | 2,643 | 191,398 | 1,840 | 2,574 | 4,732 |
| | $(1.0, 2.5]$ | 3,200,737 | 1,497 | 2,568 | 5,247 | 1,081,782 | 1,362 | 2,332 | 4,467 | 293,687 | 2,158 | 3,156 | 8,779 |
| | $(2.5, 5.0]$ | 2,186,932 | 2,270 | 4,128 | 10,568 | 246,828 | 2,117 | 3,748 | 8,313 | 338,261 | 2,533 | 4,417 | 14,638 |
| | $(5.0, 10]$ | 1,826,818 | 3,055 | 6,692 | 16,831 | 31,552 | 2,694 | 5,072 | 12,363 | 513,291 | 3,197 | 8,176 | 19,530 |
| | $(10, 25]$ | 1,681,460 | 5,225 | 15,137 | 25,749 | 805 | 3,290 | 8,142 | 17,928 | 919,982 | 5,431 | 17,092 | 27,615 |
| | $(25, 50]$ | 694,784 | 17,662 | 28,246 | 37,593 | 0 | – | – | – | 604,276 | 18,542 | 29,019 | 38,187 |
| | $(50, 100]$ | 300,996 | 33,911 | 41,749 | 48,871 | 0 | – | – | – | 297,700 | 34,049 | 41,845 | 48,941 |
| **AMR** | $(0, 0.5]$ | 8,124,067 | 407 | 1,125 | 3,186 | 634,975 | 97 | 145 | 222 | 105,375 | 1,237 | 2,684 | 10,711 |
| | $(0.5, 1.0]$ | 1,920,838 | 823 | 2,390 | 8,016 | 87,768 | 144 | 218 | 321 | 110,129 | 1,539 | 3,367 | 13,708 |
| | $(1.0, 2.5]$ | 1,861,646 | 1,117 | 3,217 | 13,840 | 32,732 | 340 | 454 | 584 | 362,381 | 1,937 | 4,488 | 17,220 |
| | $(2.5, 5.0]$ | 827,543 | 1,460 | 3,597 | 16,886 | 13,699 | 554 | 662 | 801 | 430,222 | 2,323 | 5,750 | 19,605 |
| | $(5.0, 10]$ | 784,043 | 2,076 | 4,897 | 19,281 | 6,011 | 642 | 751 | 896 | 602,870 | 2,614 | 7,301 | 21,384 |
| | $(10, 25]$ | 1,032,974 | 2,885 | 10,238 | 24,772 | 750 | 667 | 773 | 887 | 981,035 | 3,049 | 11,263 | 25,301 |
| | $(25, 50]$ | 663,726 | 4,157 | 19,824 | 33,473 | 1 | – | – | – | 662,766 | 4,166 | 19,850 | 33,488 |
| | $(50, 100]$ | 379,938 | 22,142 | 36,490 | 46,057 | 0 | – | – | – | 379,938 | 22,142 | 36,490 | 46,057 |
| **EAS** | $(0, 0.5]$ | 4,452,108 | 257 | 377 | 694 | 2,419,536 | 219 | 299 | 402 | 588,313 | 1,417 | 3,175 | 15,178 |
| | $(0.5, 1.0]$ | 830,225 | 441 | 580 | 956 | 473,651 | 400 | 496 | 620 | 127,379 | 2,677 | 7,679 | 21,953 |
| | $(1.0, 2.5]$ | 863,803 | 665 | 949 | 3,175 | 290,434 | 556 | 678 | 838 | 246,062 | 2,753 | 7,939 | 22,488 |
| | $(2.5, 5.0]$ | 566,841 | 1,124 | 2,521 | 12,564 | 59,809 | 789 | 944 | 1,167 | 295,616 | 2,849 | 8,469 | 22,505 |
| | $(5.0, 10]$ | 581,311 | 2,056 | 4,592 | 19,164 | 9,022 | 885 | 1,061 | 1,344 | 437,605 | 2,885 | 9,123 | 23,237 |
| | $(10, 25]$ | 880,951 | 2,817 | 9,193 | 24,394 | 404 | 966 | 1,205 | 1,581 | 820,256 | 3,063 | 11,130 | 25,395 |
| | $(25, 50]$ | 654,316 | 3,700 | 17,149 | 31,826 | 5 | 789 | 826 | 829 | 650,087 | 3,740 | 17,325 | 31,924 |
| | $(50, 100]$ | 469,485 | 14,392 | 32,707 | 44,104 | 1 | – | – | – | 469,398 | 14,411 | 32,711 | 44,106 |
| **EUR** | $(0, 0.5]$ | 5,501,819 | 272 | 488 | 2,002 | 952,272 | 152 | 221 | 309 | 236,433 | 2,244 | 6,848 | 21,733 |
| | $(0.5, 1.0]$ | 1,000,000 | 463 | 681 | 1,381 | 78,287 | 272 | 356 | 465 | 131,350 | 2,031 | 5,102 | 19,648 |
| | $(1.0, 2.5]$ | 1,152,780 | 752 | 1,138 | 3,489 | 11,207 | 357 | 458 | 593 | 314,046 | 2,005 | 4,688 | 18,342 |
| | $(2.5, 5.0]$ | 776,938 | 1,148 | 2,421 | 11,344 | 46 | 535 | 720 | 989 | 401,255 | 2,167 | 5,045 | 18,692 |
| | $(5.0, 10]$ | 747,466 | 1,985 | 4,451 | 18,440 | 0 | – | – | – | 566,483 | 2,500 | 6,629 | 21,036 |
| | $(10, 25]$ | 1,000,157 | 2,885 | 9,925 | 24,724 | 0 | – | – | – | 940,136 | 3,010 | 10,874 | 25,295 |
| | $(25, 50]$ | 657,062 | 3,998 | 18,937 | 32,945 | 0 | – | – | – | 654,305 | 4,011 | 19,001 | 32,986 |
| | $(50, 100]$ | 390,799 | 19,765 | 35,465 | 45,555 | 0 | – | – | – | 390,708 | 19,779 | 35,469 | 45,558 |
| **SAS** | $(0, 0.5]$ | 5,608,105 | 274 | 428 | 737 | 2,193,373 | 199 | 295 | 418 | 136,412 | 982 | 2,245 | 15,009 |
| | $(0.5, 1.0]$ | 1,481,990 | 474 | 689 | 1,188 | 668,101 | 385 | 502 | 655 | 124,372 | 1,277 | 3,114 | 15,916 |
| | $(1.0, 2.5]$ | 1,359,096 | 770 | 1,167 | 3,219 | 381,602 | 589 | 740 | 946 | 292,606 | 1,861 | 4,417 | 18,160 |
| | $(2.5, 5.0]$ | 777,344 | 1,374 | 2,839 | 12,598 | 77,540 | 892 | 1,088 | 1,427 | 389,781 | 2,337 | 5,594 | 19,259 |
| | $(5.0, 10]$ | 764,143 | 2,346 | 5,265 | 18,799 | 13,221 | 1,027 | 1,273 | 1,837 | 613,616 | 2,636 | 6,868 | 20,621 |
| | $(10, 25]$ | 1,046,826 | 3,018 | 10,510 | 24,510 | 513 | 1,135 | 1,415 | 1,929 | 1,017,345 | 3,067 | 10,952 | 24,770 |
| | $(25, 50]$ | 676,707 | 4,194 | 19,501 | 33,013 | 5 | 1,467 | 1,492 | 2,063 | 676,430 | 4,197 | 19,507 | 33,016 |
| | $(50, 100]$ | 384,158 | 22,014 | 36,446 | 46,031 | 0 | – | – | – | 384,154 | 22,015 | 36,446 | 46,031 |

### Supplementary Text

**S1 Genealogical estimation of variant age (GEVA) 14**

**S8 Inference of the ancestry shared between individuals and populations 60**

#### S1 Genealogical estimation of variant age (GEVA)

Here we introduce methodology to estimate the age of genetic variants; the point in time when a mutation gave rise to the allele observed at a particular locus in sample data. In principle, we can estimate the age of any variant segregating at any frequency in a population, without being affected by the selective forces that acted on the allele. Our method has several useful properties:

- It does not require a demographic model or assumptions about relatedness among sampled individuals. Parametric models are used within the approach to detect recombination breaks, account for error, and obtain a posterior distribution on the time to the most recent common ancestor (TMRCA) for pairs of haplotypes, but the underlying approach to estimate allele age is agnostic with respect to the genealogical process.
- It makes full use of the information available in whole genome sequencing data, combining information from both the mutation and recombination clocks inherent in population genetic data.
- It is scalable. By sampling pairs of individuals, the computational costs can be limited, with little loss of power. For example, the probability of sampling the deepest root within the coalescent tree of a population in a subsample of size $n$ is approximately $(n+1)/(n-1)$, suggesting that the most recent common ancestor (MRCA) of a subsample usually captures the MRCA of the larger sample. The probability of capturing the nearest discordant clade is more dependent on sample size (and the true age of the variant).
- It is robust to errors. Real data has sequencing or genotyping error, as well as haplotype phasing error, which can create problems in identifying haplotypes that carry a variant of interest and may create false breaks in haplotypes. We use empirically calibrated models of genotype error, which we measured in sequencing data, and we use filters to identify outliers in TMRCA distributions. The approach also makes the algorithm robust to low levels of recurrent mutation.
- It can combine information from different data sources. The core algorithm within GEVA combines information from many pairwise comparisons around a variant of interest. The comparisons can be performed across many data sets, potentially even distributed ones, with the only data needing to be shared being the parameter values of the pairwise posterior TMRCA distributions.

We refer to our method as the genealogical estimation of variant age (GEVA), which we developed as an integrated, analytical framework. We implemented GEVA in `C++` and made the source code available online.*

* https://github.com/pkalbers/geva

#### S1.1 Genealogical approach

Our goal is to estimate the age of an allele at target site $k$, of which there are $x_k$ copies in a sample of size $N$ haploid chromosomes. We assume that a mutation occurred only once at site $k$ in the history of the population and that there was no back-mutation. The allele is therefore assumed to derive from a mutation event in the genome of the common ancestor of the chromosomes that carry the allele. We assume that we know the ancestral and derived allelic states with certainty and that haplotypes have been phased.

We divide the sample into two disjoint subsets, $X_k$ and $Y_k$, consisting of carrier and non-carrier haplotypes, respectively. By tracing back the ancestry of the chromosomes in $X_k$, we expect that all of them share a common ancestor by the time of the focal mutation event and that the mutation occurred before any of them share a common ancestor with a chromosome in $Y_k$.

The genealogy of the sample at site $k$ is generally unknown. However, we can identify pairs of chromosomes whose lineages have a common ancestor either before or after the time of the focal mutation event.

- **Concordant pairs.** Any two carrier haplotypes are expected to coalesce more recently than the time of the focal mutation event. Specifically, they will have coalesced by the time of the node in the tree below where the mutation occurred.

- **Discordant pairs.** Any pair composed of one carrier and one non-carrier haplotype is expected to coalesce further back in time, prior to the time of the focal mutation event. Specifically, they will coalesce at or after the time of the node in the tree above where the mutation occurred.

Informally, the time of mutation is delimited by two time points; the time the subtree below the mutation has coalesced into a single lineage of the most recent common ancestor (MRCA) carrying the derived allele, and the time this subtree coalesced with the remaining sample. Note that we require $x_k > 1$ carrier haplotypes to form at least one concordant pair and, likewise, $x_k < N$ to form discordant pairs. In principle, however, it is possible to also estimate the age of an allele if we only find one allele copy (singletons) or if all chromosomes are fixed for the derived allele, but which we have not considered here.

#### S1.2 Sampling of haplotype pairs

The numbers of concordant and discordant pairs grow quadratically with sample size (for a fixed allele frequency), which can be computationally prohibitive. The set containing all possible concordant pairs that can be formed for a given target allele at site $k$ is given by

$$\mathcal{C}_k = \big\{\{i, j\} : \; i, j \in X_k, \; i \neq j\big\} \tag{1}$$

and the set containing all possible discordant pairs is given by

$$\mathcal{D}_k = \big\{\{i,j\} : \; i \in X_k, \; j \in Y_k\big\} \, , \tag{2}$$

which are subsets of $N(N-1)/2$ possible pairs in the sample. There are $|\mathcal{C}_k| = x_k(x_k - 1)/2$ possible concordant pairs and $|\mathcal{D}_k| = x_k(N - x_k)$ discordant pairs. We use two different sampling strategies to limit the computational cost while maintaining accuracy (described below). The maximum number of pairs sampled per group are user-defined parameters;

$$\begin{aligned} &\max\nolimits_{\mathcal{C}} && \text{for concordant pairs, and} \\ &\max\nolimits_{\mathcal{D}} && \text{for discordant pairs.} \end{aligned}$$

**Concordant pairs.** We sample concordant pairs uniformly at random. The probability that a subsample of $X_k$ includes at least one pair that spans the TMRCA for the subtree depends on the number of descendants of the two branches leading to the MRCA. In a neutral model, this partition is uniform, hence for large sample size, a random draw of $a$ chromosomes from $X_k$ will include the MRCA with probability of at least $(a-1)/(a+1)$. This implies that a random sample of concordant pairs will include the MRCA of the samples that carry the variant with high probability. Examples of the distribution of pairwise TMRCA distributions for variants at different frequencies are shown in Figure S10.

**Discordant pairs.** Our approach to sample discordant pairs is based on prioritizing non-carrier haplotypes that are the nearest genealogical neighbors to the subtree below the mutation. By the time the subsample of carrier haplotypes has merged into a single lineage, the subsample of non-carrier haplotypes will have collapsed into an unknown number of ancestral lineages, which may also collapse further before joining with the remaining lineage. Dependent on the age of the allele and the ancestral background of the sample, the number of nodes at which discordant pairs coalesce in the tree above the mutation can be relatively small; see Figure S10 for examples. A randomly formed subset of discordant pairs would likely capture a large proportion of pairs that coalesce at a node close to or at the MRCA of the sample.

To prioritize discordant pairs that are likely to coalesce more recently, we compute the Hamming distance for the set of possible pairs, measured by counting allelic mismatches between two sequences at a fixed interval to both sides relative to a given target site. Here, we scanned up to the first 5,000 positions on each side (as seen in the data). The set of pairs is sorted from low to high distance to form a priority queue. We additionally "relax" priority ranks by scanning each position in the queue, starting at the lowest distance, to remove pairs in which the same non-carrier haplotype appears more than once in consecutive order. Removed pairs are then randomly inserted at a lower rank in the queue before continuing at the next position. This is done, because the ranking of pairs based on their Hamming distance alone is limited in that the sequence interval at which pairs are compared is unlikely to be confined to the local genealogy, but rather may involve sequence variation that derived from peripheral genealogies. Also, a distinction of haplotypes predicated on carrier and non-carrier status is biased at target sites that violate model assumptions or in presence of data error. By relaxing the priority rank, we attempt to reduce the chance to include false negative non-carrier haplotypes that would otherwise be preferentially selected.

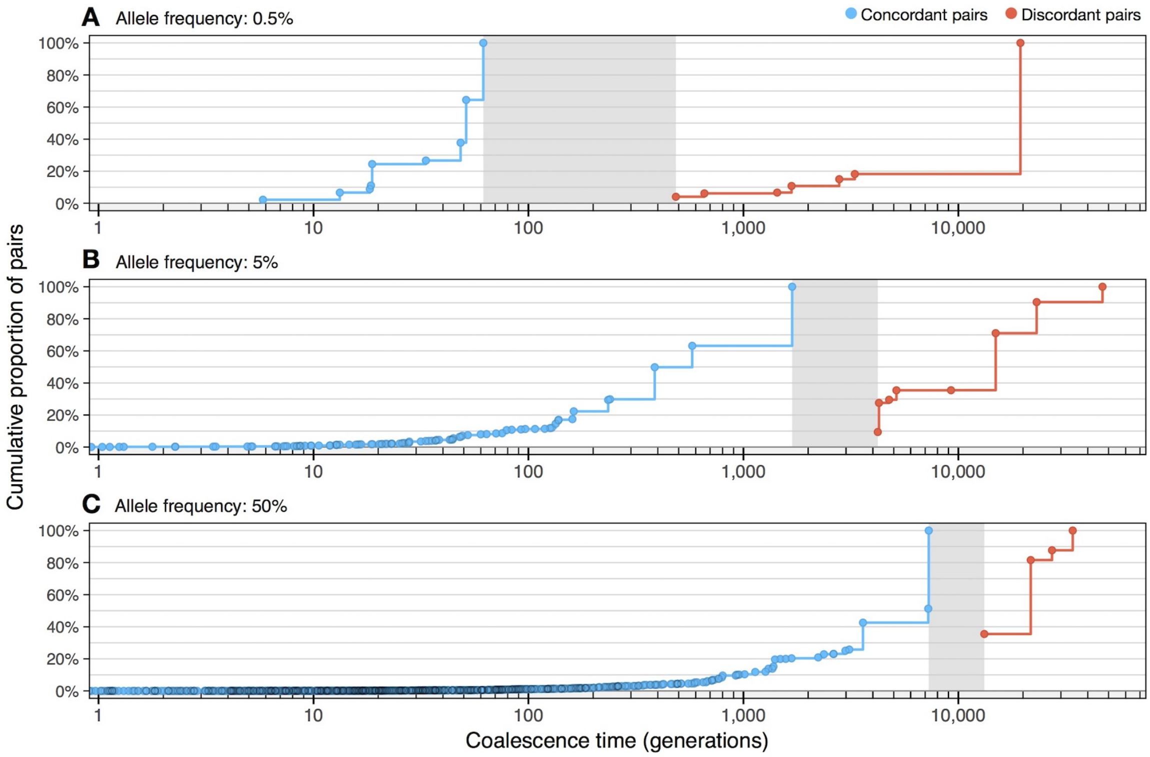

**Figure S10. Distribution of coalescence times for concordant and discordant pairs.** Coalescence time distributions for concordant and discordant pairs at three randomly selected sites in simulated data with frequencies of (A) 0.5%, (B) 5%, and (C) 50%. Data were simulated with sample size of 1,000 haplotypes, $N_e$=10,000, $\mu = 1 \times 10^{-8}$, and $r = 1 \times 10^{-8}$; see Section S4 (Script 1). Each panel shows the cumulative fraction of pairs (*y-axis*) that have coalesced back in time (*x-axis*); shown separately for pairs of concordant (*blue*) and discordant (*red*) haplotypes. Areas in *grey* indicate the branch in the underlying genealogical tree on which the focal mutation arose, delimited by the maximum and minimum of the coalescence times of concordant and discordant pairs, respectively.

#### S1.3 Inference of pairwise TMRCA

There are two main sources of information that relate to the time separating two haplotypes from their MRCA. Mutation events occur independently in each lineage and accumulate along the sequence as the ancestral haplotype is passed on over generations, and recombination events break down the length of an ancestral haplotype independently in each lineage in each generation; see schematic in Figure S11. Here we describe three coalescent-based "clock" models, which are constructed in a Bayesian setting for probabilistic inference of the TMRCA between two lineages, and where time is modeled given information about mutational differences (*mutation clock*), recombination distance (*recombination clock*), or both (*joint clock*).

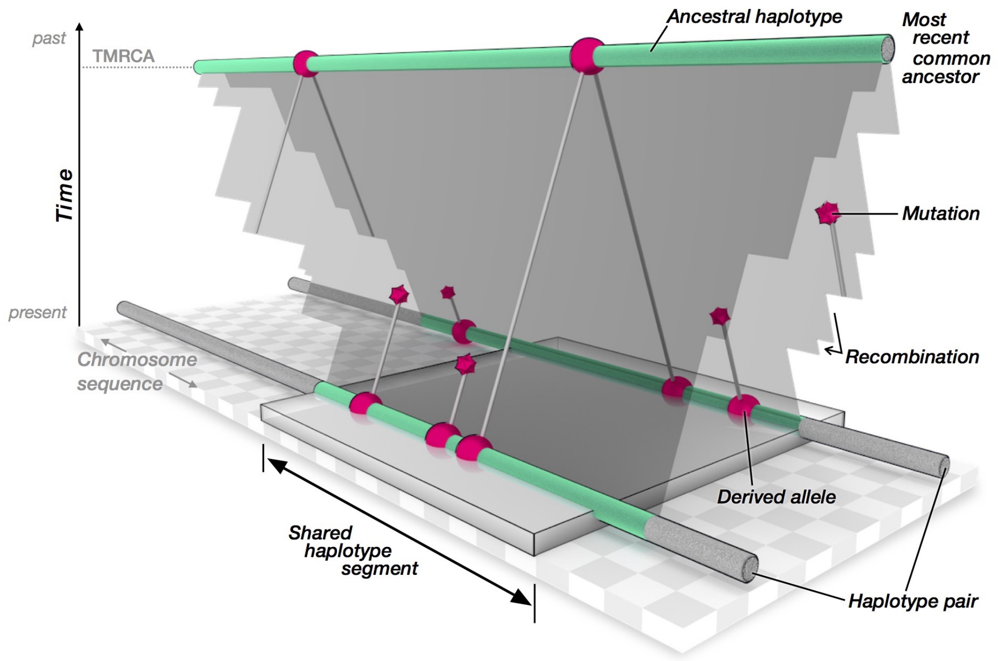

**Figure S11. Schematic of the genealogical relationship between two random haplotypes.** A random pair of haplotypes (*bottom*) share a common ancestor in the past (*top*), from whom they have inherited some piece of DNA. Over time, the ancestral haplotype sequence (*green*) has been broken down due to recombination, which occurred independently in each lineage; indicated by *cuts* in the plane connecting the two current haplotypes to their MRCA. A shared haplotype segment is locally defined as the sequence interval both haplotypes retained since inheritance from the MRCA; indicated by the *grey* block. Mutation events (*red polygons*) occurred independently in either lineage following the ancestral split from the MRCA. By using existing knowledge about the rate at which mutation and recombination events occur, it is possible to infer the time to the most recent common ancestor (TMRCA) from sequence information at the shared haplotype segment.

We define the continuous random variable $T$ for the time of coalescence, where $T = t$ takes values scaled in units of the diploid effective size of the population, $N_e$, such that

$$t = \frac{m}{2N_e} \tag{3}$$

where $m$ is the number of generations (meioses). We use the neutral coalescent to characterize the prior probability of coalescence;

$$\pi(t) \propto e^{-t}. \tag{4}$$

That is, the waiting time until the first coalescent event for two randomly sampled lineages is approximately exponentially distributed, with rate equal to 1. The choice of prior is expected to have a weak effect on the inference of TMRCA (and subsequent estimations of allele age), but depends on the order of mutation and recombination rates. For example, as the mutation rate gets large, the approach will converge on the true TMRCA irrespective of the prior. In theory, however, it would be possible to use different distributions, or estimate the prior from the data.

The setting in which each clock model operates is as follows. For a given pair of haplotypes, we treat the region they share by descent around a given focal site as known. More specifically, we assume that the genomic locations of the breakpoints that delimit the shared haplotype segment are known. We therefore assume that no recombination has occurred within the shared sequence interval in either of the two lineages considered. While this is purely theoretical, in Section S5 we propose a solution to locally infer the shared haplotype segment around a given target site, which employs a hidden Markov model (HMM) to infer the nearest breakpoints of past recombination events between two haplotype sequences.

##### S1.3.1 Mutation clock model

Following the assumptions of the infinite-sites model (ISM), mutations occur only once at each site in the history of the sample, without recurrent or back-mutations [9, 10]. It follows that the number of pairwise differences observed between two non-recombining DNA sequences is equal to the number of mutation events that occurred on both lineages since coalescence in the MRCA. The number of pairwise differences is equivalent to the number of segregating sites in a sample of two haplotypes. We further assume that all mutations in the region are observed in the data and that alleles are encoded as 0s and 1s to distinguish ancestral and derived allelic states, respectively. That is, we know the ancestral and derived states for the variant at a given locus (for example, see Section S7.1).

Given the compound mutation parameter $\theta = 4N_e\mu$, where $\mu$ is the known mutation rate per base pair per generation, mutations accumulate on each lineage independently as a Poisson process with rate $\theta/2$. We model the number of pairwise differences using the discrete random variable $S$, which follows the Poisson distribution with parameter $\theta ht$, where $t$ is the population-scaled time parameter and $h$ is the physical length of the shared haplotype segment considered, measured as the number of basepairs that make up the segment. We obtain the number of pairwise differences as the sum of allelic mismatches observed along the sequence interval. The probability to observe $S = s$ pairwise differences is given by the probability mass function (PMF) of the Poisson distribution, namely

$$P(S = s \mid \theta ht) \;=\; \frac{(\theta ht)^s}{s!} e^{-\theta ht} \,. \tag{5}$$

The likelihood function for the time parameter $t$ is proportional to the above, but requires only those terms that involve $t$ and where constant terms can be dropped, such that

$$\mathcal{L}(t \mid \theta, h, s) \;\propto\; t^s e^{-\theta ht} \tag{6}$$

from which we obtain the posterior probability of the time of coalescence as

$$\begin{aligned} p(t \mid \theta, h, s) \;&\propto\; \mathcal{L}(t \mid \theta, h, s) \;\times\; \pi(t) \\ &\propto\; t^s e^{-t(\theta h + 1)} \,. \end{aligned} \tag{7}$$

In the above, the density of the posterior probability is specified up to a missing normalising constant. The form of Equation (7) implies a Gamma distribution with shape ($\alpha$) and rate ($\beta$) parameters

$$\alpha = s + 1\,, \quad \beta = \theta h + 1 \tag{8}$$

such that the posterior density can now be written as

$$p(t \mid \theta, h, s) \;=\; \frac{(\theta h + 1)^{s+1}}{\Gamma(s+1)} t^s e^{-t(\theta h + 1)}\,. \tag{9}$$

Note that we previously defined the prior using the exponential distribution with rate equal to 1, but which is equivalent to using the Gamma distribution with $\alpha = 1$ and $\beta = 1$, such that the prior distribution is conjugate to the posterior given above. This result has been obtained previously, for example see Hein *et al.* [11, Eq. 3.45].

**Variable mutation rates.** In applications to genomic data with considerable heterogeneity of mutation rates, the model can be adjusted to consider variable rates along the genome, if such data is available. Let $\vartheta$ denote the expected value of pairwise differences over the shared haplotype segment per unit of population-scaled time. We have $\vartheta = \theta h$ if the mutation rate is uniform (as is assumed above). Otherwise, given a vector of known mutation rates per site, we compute $\vartheta = 4N_e \sum_{k=1}^{h} \mu_k$, where $\mu_k$ is the per generation mutation rate at the $k$th site in the focal nucleotide sequence of length $h$. This is used to calculate the rate of the Gamma distribution as $\beta = \vartheta + 1$.

**Conditional count of pairwise differences.** According to ISM assumptions, the number of pairwise differences observed along a non-recombinant region in two focal haplotype sequences is equal to the number of mutation events that occurred since their MRCA. However, this assumption is readily violated in applications to real (non-simulated) data; for example, due to recurrent or back-mutations, flip errors in phased haplotype data, and generally in presence of data error when alleles have been missed or falsely identified in the sequencing or genotyping process. Our model is sensitive to ISM violations, because every allelic mismatch is counted as a mutation event that separates the two focal haplotypes from their MRCA.

To account for departures from model assumptions, we exclude sites conditional on the frequency of the allele whose age we attempt to estimate. Let $f_k$ denote the frequency of the derived allele at a given target site $k$. Pairwise differences are counted by scanning along the shared haplotype region of the two sequences considered. At the $i$th site in the sequence, a mismatch is counted if $f_i \leq f_k$ or excluded otherwise. The number of pairwise differences is thereby restricted to alleles that conform to ISM assumptions; that is, mutations that occurred more recently than the focal mutation event at site $k$.

We apply this restriction to concordant pairs, as both haplotypes carry the focal allele and are expected to coalesce before the time of the focal mutation event. But it does not apply to discordant pairs, because we do not know the actual number of haplotypes in the sample that subtend the lineage at which a pair of carrier and non-carrier haplotypes join back in time. If we would restrict the count of pairwise differences in discordant pairs, inferred coalescent times are likely to be underestimated, which may likewise affect estimates of allele age. However, we expect that overestimation at discordant pairs is less problematic as it is unlikely that false allelic mismatches are equally replicated among all pairs considered. Dependent on the age of the focal allele, we may also expect that the shared haplotype segment at a discordant pair will be relatively short, as there has been more time for recombination to break down its length, thereby reducing the chance to encounter sites that violate model assumptions.

##### S1.3.2 Recombination clock model

The length of a haplotype segment shared between two sequences is delimited by two recombination events (meiotic crossovers) that occurred independently at some point in the past in either of the two lineages considered. Relative to a given target site in the genome, we characterize the surrounding haplotype segment by the two points at which the pairwise ancestral relationship changes due to recombination. To be precise, we define a *breakpoint* as the first site along the sequence that immediately follows the point at which the ancestral haplotype recombined in either of the two lineages; independently on the left and right-hand side from the target position. The full length of the focal shared haplotype is thereby enclosed by the breakpoint interval.

We use the compound recombination parameter $\rho = 4N_e r$, where $r$ is the recombination rate per site per generation. In either direction from the target site, the genetic distance to the first recombination event is exponentially distributed with parameter $\rho t$, where $t$ is the population-scaled time of coalescence. We define $D$ as a random variable for the distance along the sequence of a haploid individual. The probability to observe recombination at distance $d$ is

$$P(D = d \mid \rho t) \; = \; \rho t \, e^{-\rho t d} \tag{10}$$

and the probability that recombination occurred farther beyond along the sequence is

$$P(D > d \mid \rho t) \; = \; e^{-\rho t d} \, . \tag{11}$$

However, because recombination occurred independently along either of the two sequences considered, where only the nearest event defines a breakpoint, it follows that

$$\begin{aligned} P(D_2 = d \mid \rho t) \; &= \; 2 \times P(D = d) \times P(D > d) \\ &= \; 2\rho t \, e^{-2\rho t d} \end{aligned} \tag{12}$$

where $D_2$ denotes the breakpoint distance involving two sequences, either of which breaks first. In cases where no breakpoint is encountered before reaching the end of the chromosome, it is implied that no recombination occurred along either sequence, such that

$$P(D_2 > d \mid \rho t) \;=\; P(D > d)^2 \;=\; e^{-2\rho t d} \,. \tag{13}$$

To combine Equations (12) and (13), we can write

$$f_{D_2}(d \mid \rho t, b) \;=\; (2\rho t)^b \, e^{-2\rho t d} \tag{14}$$

where $b = 1$ if a breakpoint was found or $b = 0$ otherwise. The above can be further extended to consider the breakpoint distances on both sides, $d_L$ and $d_R$, such that the full length of the shared haplotype segment, $h$, is observed with probability

$$\begin{aligned} f_H(h) \;&=\; (2\rho t)^{b_L} \, e^{-2\rho t d_L} \;\times\; (2\rho t)^{b_R} \, e^{-2\rho t d_R} \\ &=\; (2\rho t)^{b_L + b_R} \, e^{-2\rho t h} \end{aligned} \tag{15}$$

where $b_L, b_R \in \{0, 1\}$ indicate the breakpoints on the left and right-hand side from the focal position. The likelihood function for $t$ can now be obtained from Equation (15) by ignoring multiplicative constants, such that

$$\mathcal{L}(t \mid \rho, h, b_L, b_R) \;\propto\; t^{b_L + b_R} \, e^{-2\rho h t} , \tag{16}$$

to obtain the posterior probability of coalescence time as

$$\begin{aligned} p(t \mid \rho, h, b_L, b_R) \;&\propto\; \mathcal{L}(t \mid \rho, h, b_L, b_R) \;\times\; \pi(t) \\ &\propto\; t^{b_L + b_R} \, e^{-t(2\rho h + 1)} \,. \end{aligned} \tag{17}$$

The above has the same form as the posterior probability derived for the mutation clock model; see Equation (7), Section S1.3.1. Thus, we again use the Gamma distribution, but with parameters

$$\alpha = b_L + b_R + 1 \,, \quad \beta = 2\rho h + 1 \tag{18}$$

to arrive at the formulation for the posterior density, namely

$$p(t \mid \rho, h, b_L, b_R) \;=\; \frac{(2\rho h + 1)^{b_L + b_R + 1}}{\Gamma(b_L + b_R + 1)} \, t^{b_L + b_R} \, e^{-t(2\rho h + 1)} \,. \tag{19}$$

**Variable recombination rates.** To consider recombination rate variation in our model, we can use the information provided by a high-resolution recombination map. Let $\varrho$ denote the population-scaled genetic length of the focal shared haplotype segment, such that $\varrho = \rho h$ if the recombination rate is constant over the region. The genetic length of a shared haplotype segment is taken (estimated) from the recombination map as the genetic distance between the physical positions of its breakpoints, located at sites $i$ and $j$, for which we use $f_{map}(i, j)$ as a function to return the genetic length in units of *Morgan* (M). Note that map units are usually specified in *centiMorgan* (cM), where $1\text{cM} = 0.01\text{M}$. We now can calculate $\varrho = 4N_e \times f_{map}(i, j)$, such that the rate of the Gamma distribution is $\beta = 2\varrho + 1$.

###### S1.3.3 Joint clock model

We construct a joint model that considers both mutation and recombination. The same notation is used and parameters are modeled given the assumptions (and adjustments) as described for the mutation clock (Section S1.3.1) and the recombination clock (Section S1.3.2). From there, we may immediately arrive at the joint likelihood function in support of the coalescence time $t$ as the product of the two likelihoods given in Equations (6) and (16);

$$\mathcal{L}(t \mid \theta, \rho, h, s, b_L, b_R) \;\propto\; t^{s+b_L+b_R}\; e^{-th(\theta+2\rho)} \,. \tag{20}$$

However, it is convenient to replace the term $h(\theta + 2\rho)$ with $(\vartheta + 2\varrho)$, where $\vartheta$ involves the (variable) mutation rate as described on Page 20 and $\varrho$ involves the (variable) recombination rate as described on Page 22. We therefore write

$$\mathcal{L}(t \mid \vartheta, \varrho, s, b_L, b_R) \;\propto\; t^{s+b_L+b_R}\; e^{-t(\vartheta+2\varrho)} \tag{21}$$

from which we obtain the posterior probability as

$$\begin{aligned} p(t \mid \vartheta, \varrho, s, b_L, b_R) \;&\propto\; \mathcal{L}(t \mid \vartheta, \varrho, s, b_L, b_R) \;\times\; \pi(t) \\ &\propto\; t^{s+b_L+b_R}\; e^{-t(\vartheta+2\varrho+1)} \,. \end{aligned} \tag{22}$$

Conveniently, both the mutation and recombination clock models specify the Gamma distribution for the calculation of the posterior density. We therefore have

$$\begin{aligned} \alpha \;&=\; s + b_L + b_R + 1 \,, \\ \beta \;&=\; h(\theta + \rho) + 1 \;=\; \vartheta + 2\varrho + 1 \end{aligned} \tag{23}$$

to calculate the posterior density as

$$p(t \mid \vartheta, \varrho, s, b_L, b_R) \;=\; \frac{(\vartheta + 2\varrho + 1)^{s+b_L+b_R+1}}{\Gamma(s + b_L + b_R + 1)} t^{s+b_L+b_R} e^{-t(\vartheta+2\varrho+1)} \,. \tag{24}$$

##### S1.4 Composite posterior estimation of variant age

Our approach to estimate allele age is similar to existing composite likelihood methods that are applied to solve problems where the full likelihood function is unknown or intractable. Here, coalescence between a haplotype pair (concordant or discordant) is seen as a lower-dimensional feature of the local genealogical structure of the sample at a given focal variant. We combine information from hundreds or thousands of pairs to obtain an estimate of the time the allele has emerged through mutation.

The posterior density of the time to coalescence for a given pair is defined for the clock models presented above (Section S1.3). Each clock model calculates the posterior using the Gamma distribution with parameters $\alpha$ and $\beta$. For simplicity, we express the posterior density using $p(t \mid \lambda)$, where we use $\lambda = \{\alpha, \beta\}$ to connote the parameters determined from haplotype data under a specific model. The probability of coalescence more recently than (or at) time $t$ is obtained from the cumulative distribution function (CDF) of the posterior density;

$$\overline{\Lambda}(t \mid \lambda) \;=\; P(T \leq t \mid \lambda) \;=\; \int_0^t p(u \mid \lambda)\, du\,. \tag{25}$$

The probability of coalescence subsequent to time $t$ (i.e. further back in time) is likewise

$$\begin{aligned}\underline{\Lambda}(t \mid \lambda) \;=\; P(T > t \mid \lambda) \;&=\; \int_t^\infty p(u \mid \lambda)\, du \\ &=\; 1 - \overline{\Lambda}(t \mid \lambda)\,.\end{aligned} \tag{26}$$

We use the notation $\{i, j\} \leftarrow \mathcal{C}_k$ to indicate that a pair was sampled (without replacement) from the set of possible concordant pairs and, likewise, $\{i, j\} \leftarrow \mathcal{D}_k$ for discordant pairs, where $i, j$ indicate the two haplotypes in a pair. Sampling from either set is done as described in Section S1.2, but where we additionally remove concordant and discordant pairs, before sampling, that appear to be inconsistent, for which we employ a heuristic algorithm (described in Section S1.5).

The age of an allele observed at target site $k$ is estimated from the composite posterior distribution;

$$\Phi_k^\tau(t) \;\propto\; \prod_{\{a,b\} \leftarrow \mathcal{C}_k} \overline{\Lambda}\big(t \mid \lambda_k^\tau(a,b)\big) \;\times\; \prod_{\{c,d\} \leftarrow \mathcal{D}_k} \underline{\Lambda}\big(t \mid \lambda_k^\tau(c,d)\big) \tag{27}$$

where $\tau$ indicates the coalescent clock model used. The composite posterior distribution can now be obtained over $t \in (0, \infty)$, from which we take the mode as a point estimate of variant age.

#### S1.5 Heuristic method to reject outlier pairs

There are several sources of error that may adversely affect the estimation of allele age when using the composite posterior approach described above. For example, a given focal variant may have been falsely called or missed, the allele may have been lost due to back-mutation, or some of the shared haplotype segments may have been inferred incorrectly. To reduce the impact of such outliers on the estimation process, we perform quality control on the set of pairs before they are included in the computation of the composite posterior distribution.

Using a simple, heuristic algorithm, a time threshold is ascertained above and below which concordant and discordant pairs are rejected, respectively; see Figure S12. The number of available concordant pairs is given by $n_{\mathcal{C}}$, where $1 \leq n_{\mathcal{C}} \leq \max_{\mathcal{C}}$, and the number of discordant pairs is given by $n_{\mathcal{D}}$, where $1 \leq n_{\mathcal{D}} \leq \max_{\mathcal{D}}$. The mean of the inferred coalescence time distribution is taken as a point estimate for the TMRCA of each pair. It follows from the

Gamma distribution that the mean is $\mathrm{E}[T] = \alpha/\beta$, where $\alpha, \beta$ are determined from haplotype data per pair as defined for a given clock model. The sets of concordant and discordant pairs are sorted, separately, from lowest to highest mean TMRCA. For simplicity, we have

$$\bar{c}_1,\ \bar{c}_2,\ \bar{c}_3,\ \ldots,\ \bar{c}_{n_\mathcal{C}} \ \text{ and } \ \bar{d}_1,\ \bar{d}_2,\ \bar{d}_3,\ \ldots,\ \bar{d}_{n_\mathcal{D}}$$

where $\bar{c}_i$ and $\bar{d}_i$ connote the mean TMRCA of a given concordant and discordant pair, respectively. We find a threshold to reject pairs if $\bar{d}_1 < \bar{c}_{n_\mathcal{C}}$, or we reject none otherwise. The threshold is placed such that the minimum total number of pairs is rejected. However, we keep the most recent concordant pair and the oldest discordant pair, so as to ensure that there is at least one pair in each group.

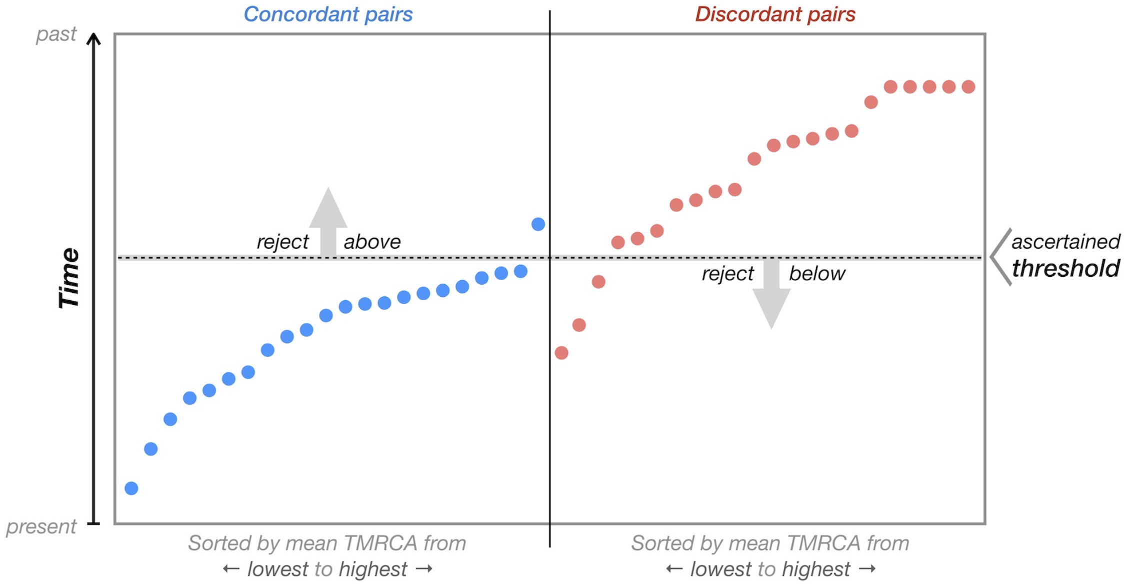

**Figure S12. Schematic concordant and discordant pair filtering.** Available concordant (*left*) and discordant pairs (*right*) are independently sorted by mean TMRCA. Pairs are rejected if distributions overlap. We use a heuristic algorithm to ascertain a threshold above and below which concordant and discordant pairs are rejected, respectively. The threshold is determined such that the minimum total number of pairs is rejected.

##### S1.5.1 Quality score

The proportion of concordant and discordant pairs rejected for a variant may provide a simple metric to evaluate the quality of its age estimate (resulting from a given clock model). We calculate a quality score, $QS$, as

$$QS = 1 - \max\left\{\frac{r_\mathcal{C}}{n_\mathcal{C}}, \frac{r_\mathcal{D}}{n_\mathcal{D}}\right\} \tag{28}$$

where $r_\mathcal{C}$ is the number of rejected concordant pairs and $r_\mathcal{D}$ the number of rejected discordant pairs. Values near or equal 1 indicate high quality, and values near 0 indicate low quality; note that at least one pair is retained in each group, such that $0 < QS \leq 1$.

#### S2 Shared ancestry inference

The ancestry shared between two haploid genomes can be described by the cumulative coalescent function (CCF), which is a monotonically increasing "coalescent profile" that expresses the fraction of a given (*target*) genome that has coalesced with another (*comparator*) genome up to a given point back in time. We use a dynamic programming technique for maximum likelihood decoding of the CCF, which operates on the distribution of allele sharing observed between two haplotype sequences, given the (estimated) ages of alleles. The method assumes independence of variants and ignores error in age estimates. We implemented this method in C++ and made the source code available online.* The rationale of the method is illustrated in Figure S13; the following section describes the algorithm in detail.

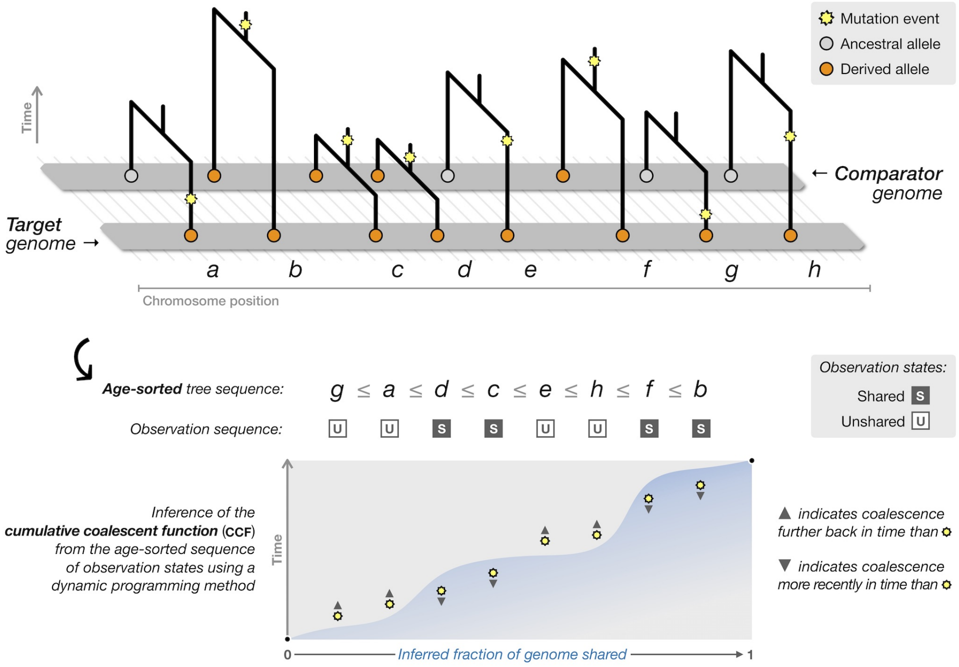

**Figure S13. Schematic of shared ancestry inference using dynamic programming.** The figure shows a schematic representation of two genomic sequences (*top*); labels $a, b, \ldots, h$ identify the derived alleles (*orange*) carried by a given target genome, for which allele age information is available. The genealogical relationship between the target and a given comparator genome is characterized by a sequence of trees. The mutation event that gave rise to an allele is indicated (*yellow*) on each tree at some point in the past. We define two observation states; *shared* (both the target and comparator carry the derived allele) and *unshared* (only the target carries the derived allele). Unshared alleles where only the comparator carries the derived allele are ignored. Given knowledge about the age of alleles, we sort the trees by the time of mutation (from youngest to oldest) to obtain the age-sorted observation sequence of shared and unshared states (*middle*). The time of coalescence between the two genomes considered is indicated by the shared or unshared state (*bottom*); we expect shared alleles to coalesce more recently than the time of mutation, and unshared alleles to coalesce further back in time. Implicitly, this involves the cumulative distribution $P(T_2 \leq t)$ of the probability that the time of coalescence between the two lineages ($T_2$) is not larger than the time of mutation ($t$) at each shared site, and conversely $P(T_2 > t)$ at each unshared site, while assuming that mutations and the trees on which they occurred are independent. Knowledge about the times of the mutations carried by a given target genome allows us to determine the relative order of events of lineages coalescing with the target genome. We thereby seek to infer the fraction the target genome shares with the comparator genome from the sequence of coalescent events as a function of time; referred to as the cumulative coalescent function (CCF), which we infer using a dynamic programming method. In the figure, the inferred CCF is indicated by the shaded area (*blue*).

* https://github.com/pkalbers/ccf

##### S2.1 Cumulative coalescent function (CCF)

Let $M$ be the number of mutations carried by a given target genome for which the age, denoted by $\phi$, has been estimated. These are sorted from most recent to oldest age to obtain a sequence of time points, indexed by $i = 0, 1, 2, \ldots, M-1$, such that

$$\phi_0 \leq \phi_1 \leq \phi_2 \leq \ldots \leq \phi_{M-1}.$$

Corresponding to the above, we generate a sequence of observations,

$$\omega_0,\ \omega_1,\ \omega_2,\ \ldots,\ \omega_{M-1}$$

where each observation in the age-sorted sequence is encoded as

$$\omega_i = \begin{cases} 1 & \text{if } \textit{shared} \\ 0 & \text{if } \textit{unshared} \end{cases} \tag{29}$$

to indicate the two possible observation states; *shared* if both the target and the comparator genomes carry the mutation, or *unshared* if only the target genome carries the mutation. The shared state indicates that the two genomes coalesce more recently than (or at) the time of the mutation event, while the unshared state indicates coalescence further back in time. Note that we only consider mutations carried by the target genome, because the times of unshared mutations carried by only the comparator are independent from the timings of coalescent events along the target genome.

The fraction of the ancestry a given target genome shares with another genome is discretized at the level of a single nucleotide. This being impractical for analytical purposes, we use a discrete choice approach as an approximation technique. We define $\{\delta_j\}_{j=0\ldots S}$ as the parameter space of possible states for the CCF, where $0 < \delta_j \leq 1$. For $S$ states, we obtain

$$\delta_j = \frac{j}{S}, \text{ for } j = 1, 2, \ldots, S \tag{30}$$

and set $\delta_0 = \varepsilon$, where $\varepsilon$ is reasonably small; $0 < \varepsilon \ll 1/S$.

The steps of the dynamic programming algorithm for inference of the CCF are as follows.

**Initialization.** Let $\mathbf{A}, \mathbf{B}$ denote two matrices of size $M \times (S+1)$. These are initialized at the first site in the sequence, for states in $j = 0, 1, 2, \ldots, S$,

$$\mathbf{A}_{0,j} = \omega_0 \delta_j \ + \ (1 - \omega_0)(1 - \delta_j) \tag{31}$$

$$\mathbf{B}_{0,j} = 0. \tag{32}$$

Implicitly, $\mathbf{A}_0(j \mid \omega_0)$ is equivalent to the binomial likelihood function over the discretized parameter space $\{\delta_j\}_{j=0\ldots S}$, given the observation $\omega_0$. Matrix $\mathbf{B}$ stores the maximum likelihood estimate at each state in the subsequent recursion step.

**Recursion.** Moving along the sequence of observations, for $i = 0, 1, 2, \ldots, M-1$, we compute for each state in $j = 0, 1, 2, \ldots, S$,

$$\mathbf{A}_{i,j} = \big(\omega_i \delta_j \ + \ (1-\omega_i)(1-\delta_j)\big) \times \max_{k=0\ldots j} \big[\mathbf{A}_{i-1,k}\big] \tag{33}$$

$$\mathbf{B}_{i,j} = \operatorname*{argmax}_{k=0\ldots j} \big[\mathbf{A}_{i-1,k}\big]. \tag{34}$$

The recurrence relation in $\mathbf{A}_{i,j}$ is to update the likelihood incrementally, where $\mathbf{B}_{i,j}$ stores the state that makes $\mathbf{A}_{i,j}$ most likely. Note that $k$ may not exceed the current index $j$, which is to ensure that transitions to the current state from a higher state have zero probability.

**Termination.** We define **Z** as a zero-indexed array of length $M$ to store the index of states in the subsequent traceback step. Likewise, we define **P** as a zero-indexed array of length $M$, in which the inferred path sequence will be stored. At the last position in both arrays, we set

$$\mathbf{Z}(M-1) = S \tag{35}$$

$$\mathbf{P}(M-1) = 1 \tag{36}$$

which reflects the underlying assumption that the ancestry of the two genomes can be traced back, eventually, to a single origin in the past.

**Traceback.** We seek to find the optimal path through the parameter space $\{\delta_j\}_{j=0\ldots S}$ that maximizes the likelihood given the observation sequence. Tracing back from the terminal position, for $i = M-1,\ M-2,\ \ldots,\ 1$, we compute

$$\mathbf{Z}(i-1) = \mathbf{B}_{i,\mathbf{Z}(i)} \tag{37}$$

$$\mathbf{P}(i-1) = \delta_{\mathbf{Z}(i-1)}. \tag{38}$$

We then obtain the CCF by mapping the inferred state path at $\mathbf{P}_i$ to the corresponding variant age $\phi_i$, which gives the inferred fraction of the genome shared at each time point in $\{\phi_i\}_{i=0\ldots M}$.

Finally, we define the CCF as a function of time, denoted by $\Lambda_t$ for $t \in (0, \infty)$, where $\Lambda_0 = 0$ and $\Lambda_\infty = 1$. In practise, we approximate $\Lambda_t$ using linear interpolation over a fixed grid of time points; for example, to jointly assess the CCFs from multiple individuals at the same time points in downstream analyses. For this, we write

$$\Lambda_t \ = \ \frac{\mathbf{P}_{k-1}\,(\phi_k - t) + \mathbf{P}_k\,(t - \phi_{k-1})}{\phi_k - \phi_{k-1}}\ , \ \text{ for } \ k = \operatorname*{argmin}_{i=1\ldots M} \big[\ |\phi_i - t|\ \big] \tag{39}$$

where $t$ is taken from a fixed grid of $L$ time points in $\{t_i\}_{i=1\ldots L}$, which may or may not overlap with the estimated allele ages in $\{\phi_i\}_{i=0\ldots M}$.

##### S2.2 Coalescent intensity function (CIF)

For a given time interval (or "epoch"), we infer the intensity of coalescence between a target and comparator genome from the rate of change (gradient) of the CCF. Specifically, we compute the coalescent intensity function (CIF) as

$$\lambda(t_i, t_j) \;=\; \log\left[1 - \Lambda(t_i)\right] - \log\left[1 - \Lambda(t_j)\right] \tag{40}$$

where $t_i < t_j$, which denote the lower and upper time points that delimit a given epoch, respectively. The logarithmic difference approximates the percent change as a smooth function over time, which is appropriate given the continuous-time coalescent process. Note that the above is only defined for $\Lambda_t < 1$.

The CCF approximates the cumulative distribution of the time until the comparator genome has fully coalesced with the target genome, where the fraction shared between the two genomes is defined proportionally to time $T$ as a random variable. Equivalently,

$$\Lambda_t \;\equiv\; F(T \leq t) \;\approx\; 1 - e^{-\lambda(t)\; t} \tag{41}$$

which follows from the assumption that coalescent events are mutually independent and the time to coalescence is approximately exponentially distributed with (instantaneous) rate $\lambda(t)$. Note that $t$ is scaled in units of constant size $N_0$ (or, more commonly, denoted by $N_e$). The expression given in Equation (40) is therefore equivalent to the intensity parameter of a Poisson process; namely

$$\lambda(t_i, t_j) \;\equiv\; \int_{t_i}^{t_j} \lambda(u)\, du \;=\; \bar{\lambda} \times (t_j - t_i) \tag{42}$$

where $\bar{\lambda}$ is the average (constant) rate of coalescence during the epoch considered.

##### S2.3 Effective population size ($N_e$) equivalent

Generally, changes in population size induce changes in the rate of coalescence, such that the inverse of the coalescent rate is equal to the relative population size at time $t$. Thus, the time-variable population size is given by the relation $N_t = N_0/\lambda_t$, where $N_0$ is the size constant by which time is scaled. Note that we have $\lambda_t = 1$ if population size is constant over time (equal to $N_0$).

Here, we use the CIF inferred for a given target genome to estimate the size of the ancestral population from which it derived in the past. Because the ancestry of a single genome is a distribution over many genealogical relationships, tracing back to different populations and at different points in time, we consider the pairwise coalescent history inferred between the target genome and a larger sample of comparator genomes, to obtain an estimate of the ancestral population size from the strongest signal of shared ancestry within a given time interval. Analogous to notation of the effective population size in population genetics modeling, we refer to this estimate as the $N_e$ *equivalent* (denoted by $N_{\tilde{e}}$ below).

For a given epoch that is delimited by time points $t_i, t_j$, we compute

$$N_{\tilde{e}}(t_i, t_j) \;=\; N_0 \times (t_j - t_i) \times \max_{k=1\ldots n} \Big[\lambda_k(t_i, t_j)\Big]^{-1} \qquad (43)$$

where $k$ identifies the CIF in a sample of $n$ comparators available for a given target. Note that we use the maximum CIF to obtain a non-parametric estimate of the population size, but which is restricted by the empirically determined parameter space $\{\lambda_k\}_{k=1\ldots n}$ of the sample. Hence, in practice, CIFs are inferred against comparators from a large sample of individuals with both similar and diverse ancestral backgrounds, as $N_{\tilde{e}}$ tends to be overestimated if the intensity of coalescence with available comparators is low. Likewise, the length of an epoch is chosen to allow sufficiently many comparators to coalesce with a given target.

##### S2.4 Aggregation of CCFs across chromosomes

The dynamic programing algorithm presented in Section S2.1 enables us to rapidly compute the CCF between every pair of haploid chromosomal sequences in large sample data sets. To generate a summary of the ancestry shared between diploid individuals (or groups of individuals), we combine information across the different chromosomes as follows. Consider a sample of $N$ diploid individuals whose genome is constituted of $K$ chromosomes (for example, $K = 22$ if considering autosomes in humans). We infer the CCF separately for each chromosome $k$, between each of the $2N$ target sequences in turn against $2N - 1$ comparator sequences. CCFs are approximated at a fixed grid of $L$ time points to subsequently match chromosomes $1, 2, \ldots, K$ at every time point in $\{t_i\}_{i=1\ldots L}$. Profiles are aggregated, first, among diploid individuals and per chromosome $k$, by computing the average fraction $\Lambda_t$ at each time point $t$, and then across chromosomes by computing the (weighted) mean, where weights might be assigned to different chromosomes, for example, conditional on the number of dated variants available per chromosome. This process is illustrated in Figure S14, but also explained in more detail below.

There are four CCFs inferred between the two haploid sequences of a diploid target individual $I$ and the two haploid sequences of comparator individual $J$, for $I \neq J$; namely

$$\Lambda_t^k(I_0, J_0),\; \Lambda_t^k(I_0, J_1),\; \Lambda_t^k(I_1, J_0),\; \Lambda_t^k(I_1, J_1)$$

where $k$ refers to the chromosome currently considered, and the subscripts $0, 1$ indicate the haploid sequence per diploid individual. We also obtain two CCFs between the haploid sequences of the same individual ($I = J$); namely

$$\Lambda_t^k(I_0, I_1),\; \Lambda_t^k(I_1, I_0).$$

The aggregated CCF between the target and the comparator individual is denoted by $\Lambda_t^*(I,J)$, which we compute as

$$\Lambda_t^*(I,J) = \sum_{k=1}^{K} \left( \frac{1}{4} \sum_{a=0}^{1} \sum_{b=0}^{1} \Lambda_t^k(I_a, J_b) \right) \times W_k \tag{44}$$

where $W_k$ refers to the weight applied to each chromosome, for example, calculated as $W_k = v_k/u$, where $v_k$ is the number of variants dated on chromosome $k$, and $u$ is the sum of variants dated across all chromosomes considered. Similarly, to aggregate the CCFs within the same individual, we compute

$$\Lambda_t^*(I) = \sum_{k=1}^{K} \frac{1}{2} \left( \Lambda_t^k(I_0, I_1) + \Lambda_t^k(I_1, I_0) \right) \times W_k \,. \tag{45}$$

The above can be extended to aggregate the CCFs obtained for the haploid sequences of species with higher ploidy, or across groups of individuals to summarize the ancestry shared between defined demographic units.

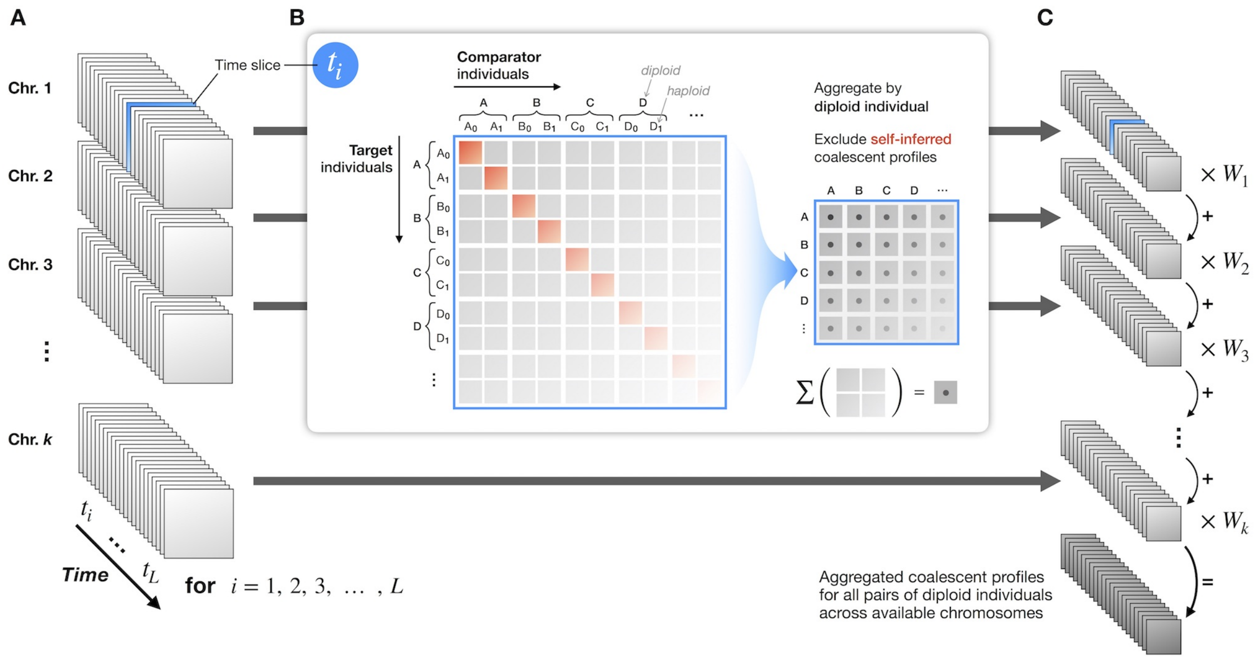

**Figure S14. Aggregation of coalescent profiles per diploid individual.** Panel **A** shows the schematic representation of the data available after inference of the CCF between each pair of haplotypes in a large sample of $N$ diploid individuals. Each chromosome $1, 2, \ldots, K$ can be represented as a three-dimensinoal matrix of size $2N \times 2N \times L$, where $L$ is the number of time points to which CCFs have been approximated. Panel **B** illustrates the aggregation of CCF data per diploid individual. Every time "slice" in $\{t_i\}_{1\ldots L}$ is represented as a $2N \times 2N$ matrix containing the inferred fraction ($\Lambda_i$) each of the $2N$ target haplotypes shares with each of the $2N - 1$ comparator haplotypes. We compute the average fraction among the four data points per pair of diploid individuals. Panel **C** shows the resulting, averaged matrices for each chromosome $k$, which are further aggregated into a single three-dimensional matrix of size $N \times N \times L$. This is done by computing the weighted mean across chromosomes, where weights ($\sum W_k = 1$) are assigned, for example, to up or down weight chromosomes with higher or lower numbers of dated variants, respectively.

#### S3 Assessment of genotype error in genomic data

In this section, we describe the construction of an empirical genotype error profile. Information derived from the error profile was used to

- reproduce realistic rates of error in simulated data (described in Section S4),
- develop a method for detecting shared haplotype segments while being robust towards data error when applied to real genomic data sets (Section S5), and
- evaluate the performance of GEVA (which includes the method for shared haplotype detection) and characterise the effects on age estimation in simulations of large sample data before and after the inclusion of error (Section S6).

Any assessment of error in empirical data requires the existence of an error-free "gold standard" against which data can be compared; provided that data were obtained on the same biological sample. We used genotype data from the Illumina Platinum Genomes (IPG) project as a reference "truthset" [4],* against which we assessed error in matched data from the 1000 Genomes Project (TGP) [3];† for human assembly GRCh37 (hg19), which are available for both panels.

In the following, we use $g$ to denote the assumed true genotypic state (as seen in IPG), and $\tilde{g}$ to denote the genotype that has been observed in the assessed data set (TGP), at a given locus and for the same individual. We define the coefficient $e_{i \to j}$ to denote the rate at which a true genotype $g_i$ was observed as genotype $\tilde{g}_j$, where the subscripts $i, j \in \{0, 1, 2\}$ indicate the genotypic state; homozygous for the reference allele (0), heterozygous (1), or homozygous for the alternative allele (2).

##### S3.1 Data preparation

The TPG sample comprises 2,504 individuals sequenced (or genotyped) at >80 million SNPs, for which data has been generated using a combination of low-coverage whole-genome sequencing ($> 4\times$), high-coverage exome sequencing ($> 50\times$), and microarray genotyping [3]. Data from IPG comprises 4.7 million SNPs generated using high-coverage whole-genome sequencing of a 17-member, three-generation family of European ancestry (`CEPH 1463`); namely, four individuals in the founder generation, two in the parental generation, and eleven children. The IPG sample has been sequenced at $50\times$ coverage on Illumina HiSeq 2000 and variants have been called in accordance with different methods to resolve conflicts among different call sets. The two parents (IDs `NA12877` and `NA12878`) have been additionally sequenced at $200\times$ coverage and variant data has been validated based on Mendelian inheritance constraints from pedigree information [4].

Cell lines from `CEPH 1463` are a well-characterized model system and have been sequenced or genotyped in several studies. Data for individual `NA12878` was available in both the IPG

* `https://emea.illumina.com/platinumgenomes.html`
† `ftp://ftp.1000genomes.ebi.ac.uk//vol1/ftp/release/20130502/`

and TGP panels. We compared genotype data that we extracted from both panels for this individual, where the genotypic state observed in IPG was assumed to be the "true" state. Although the possibility that IPG retained misclassified genotypes at a certain fraction of variant sites cannot be excluded, we assumed this fraction to be negligibly small. Likewise, the impact of other sources of error, such as somatic mutations that occurred in the sampled biological material and thereby may produce different results when processed on different platforms, were assumed to be negligible.

We extracted genotype data for all autosomes and matched sites by chromosome position between the two panels. We only considered biallelic SNPs, and we excluded sites at which the reference or alternate alleles were inconsistent between IPG and TGP. Data were additionally filtered using the accessibility mask available from IPG, thereby retaining only sites called with high confidence in IPG. Conversely, however, we did not filter sites based on high-confidence regions available for the TGP panel, due to the underlying intention to measure error as typically encountered in analyses of genomic data. The matching processes is illustrated in Figure S15.

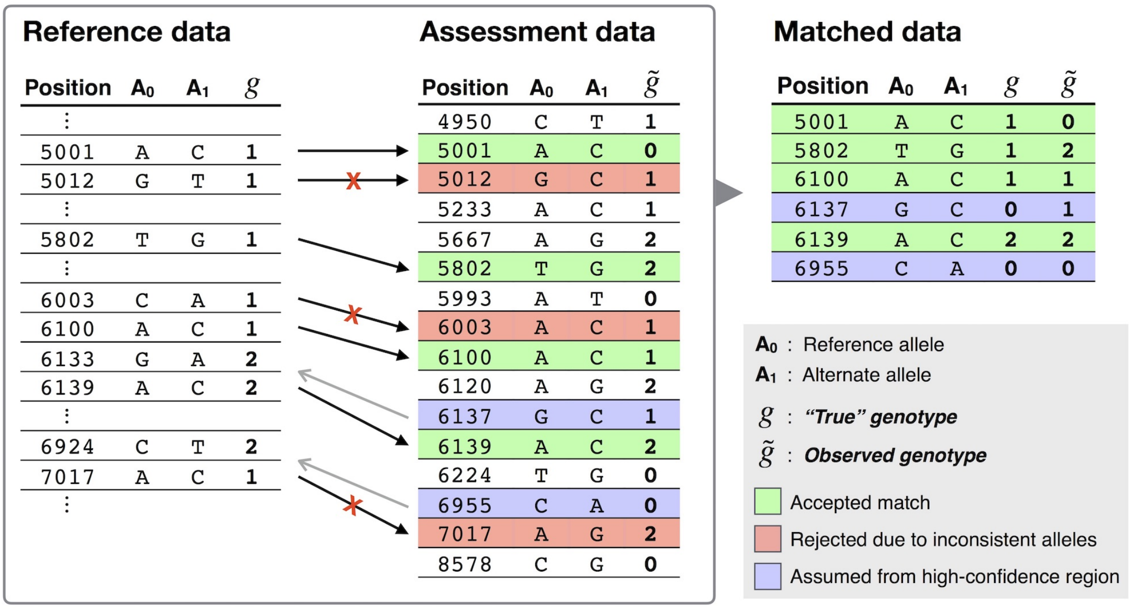

**Figure S15. Schematic of the genotype matching process.** Genotype information of individual NA12878 were extracted from the Illumina Platinum Genomes (IPG) truthset and data from the 1000 Genomes Project (TGP) Phase 3, from all autosomes, and sites were matched by chromosome position (GRCh37). We used the accessibility mask available for the IPG panel to retain variant sites called with high-confidence (indicated by gaps in the figure). Sites were removed if reference or alternate alleles did not match between IPG and TGP. Because the reference truthset (IPG) did not contain genotypes homozygous for the reference allele, these were assumed from high-confidence regions if present in the assessment dataset (TPG); indicated by left-pointing arrows. Figure modified from [12].

Note that available IPG data did not contain variants called as being homozygous for the reference allele ($g_0$). This is because the high-confidence regions in IPG have been identified in the individual call sets by collating sites that were called as being homozygous for the reference allele and monomorphic in the sample [4]. Variants homozygous for the alternate allele ($g_2$) have not been removed. Here, we made the following assumption to be able to measure error proportions pertaining to each genotypic state. The true state was assumed to be of the $g_0$ type if the position of a variant site in TGP was within high-confidence regions of the IPG accessibility mask. We assumed that high-confidence intervals encompassed variants which would have been reported as a different type otherwise.

#### S3.2 Genotype error profile

We measured genotype error at 76 million comparisons between IPG and TGP. Of those, 73.2 million were homozygous for the reference allele. While the vast majority of genotypes (>99%) have been called (or typed) without error, we found 0.08% of genotypes to be misclassified. The confusion matrix given in Table S3 shows the overall proportion of states observed per true genotypic state. We found that the density of erroneous genotypes increased towards the telomeric and centromeric regions on each chromosome (Figure S16A), where we see error densities of >0.2% on all chromosomes, but >1% on most chromosomes, on a genome-wide background of ~0.1% on average.

**Table S3. Relative error rates measured per genotype class.** Genotype data from individual NA12878 was available at 76 million comparisons between the Illumina Platinum Genomes (IPG) truthset and the 1000 Genomes Project (TGP) across autosomes. Error rates were calculated as the relative proportion a given true genotype $g_i$ was observed as $\tilde{g}_j$, where $i, j \in \{0, 1, 2\}$; thus, summing to 100% per column. The total number of genotypes assessed per true genotype class is given below.

| Observed genotype | True genotype | | |
|---|---|---|---|
| | $g_0$ | $g_1$ | $g_2$ |
| $\tilde{g}_0$ | 99.942% | 0.550% | 0.034% |
| $\tilde{g}_1$ | 0.041% | 99.282% | 0.231% |
| $\tilde{g}_2$ | 0.017% | 0.168% | 99.735% |
| *Total* | 73,211,532 | 2,076,098 | 1,326,955 |

Next, we constructed a frequency-dependent genotype error profile. Assessed sites were assigned their population frequency as observed in the TGP sample and then pooled into 200 evenly distributed frequency bins. We calculated

$$e_{i \to j} = P(g_i \to \tilde{g}_j | f), \text{ for } i, j \in \{0, 1, 2\} \quad (46)$$

as the relative rate a true genotype $g_i$ was observed as $\tilde{g}_j$ for sites at allele frequency $f$ (recorded at the mean per bin), and normalized to sum to 1 per true genotype class; $\sum_{j=0}^{2} e_{i \to j} = 1$. Results are shown in Figure S16B.

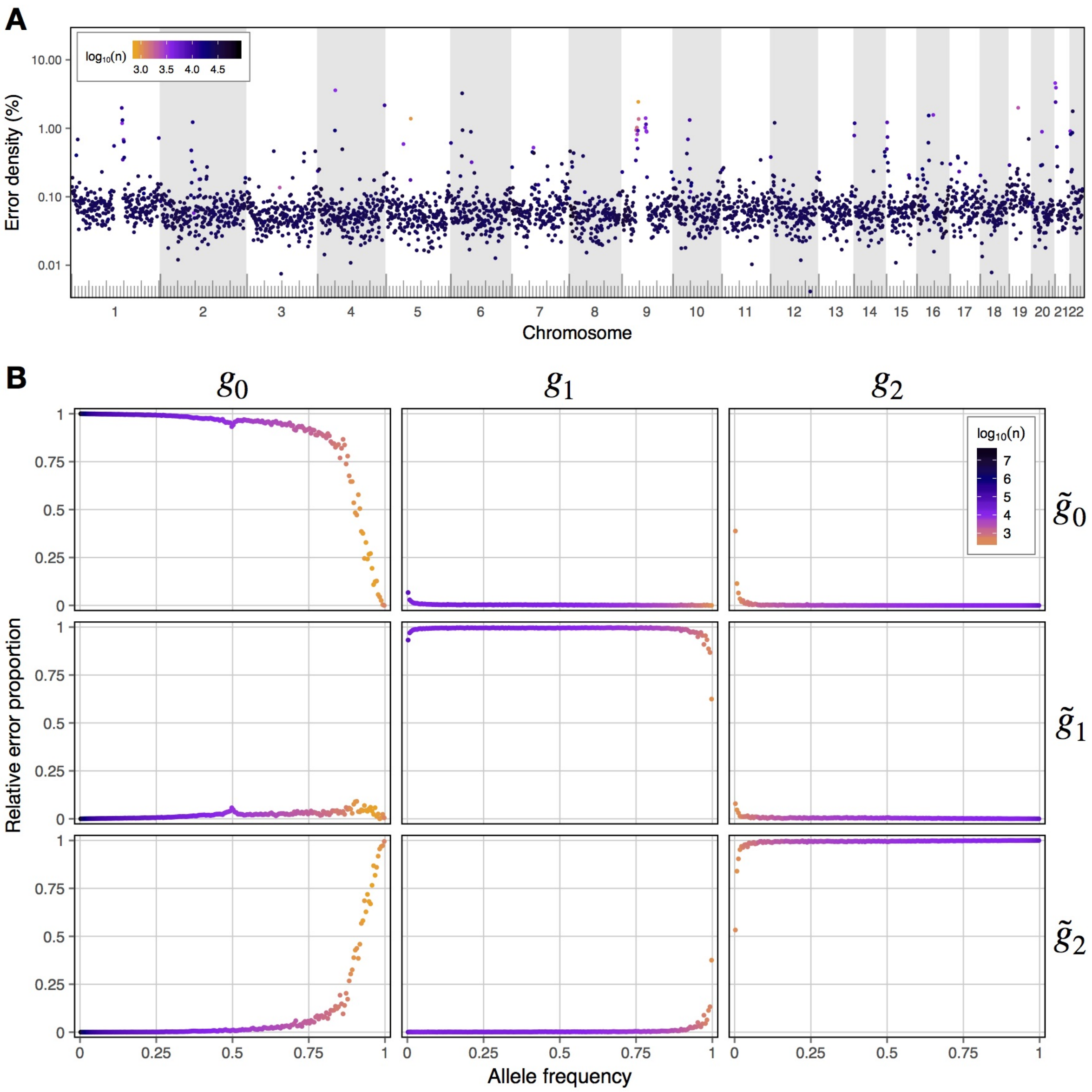

**Figure S16. Genotype error measured in data from the 1000 Genomes Project.** Genotype error was measured at 76 million comparisons between the Illumina Platinum Genomes (IPG) truthset and the 1000 Genomes Project (TGP), on genotype data extracted for the same individual (`NA12878`) from all autosomes. Panel **A** shows measured error densities by region on Chromosomes 1-22. The density of misclassified genotypes ($g \neq \tilde{g}$) was calculated in equally sized chunks of 1 Mb size along the length of each chromosome. Error density was calculated as the number of misclassified genotypes divided by the total number of genotypes per chunk; percentage shown on log-scale. Colors indicate the number of genotypes per chunk (see legend). The ruler at the bottom indicates the physical length per chromosome, where longer tick marks sit 10 Mb apart. Panel **B** shows the confusion matrix of relative error measured per genotypic class, given the allele frequency at matched sites in the TGP sample. The set of matched genotypes was pooled into 200 equally sized allele frequency bins. Relative error, $e_{i \to j}$, was calculated by counting $g_i$ observed as $\tilde{g}_j$ and dividing by the sum of $g_i$ per frequency bin; for $i, j \in \{0, 1, 2\}$. Colors indicate the number of genotypes per bin (see legend).

#### S4 Generation of simulated data

We performed coalescent simulations using `msprime` software [13]. The software stores the history of the simulated sample, which can be queried in downstream analyses through its `python` interface. We used data generated from two main simulations (described below); in the following referred to as data sets $\mathcal{A}$ and $\mathcal{B}$. We further modified haplotype data in a copy of data set $\mathcal{B}$ to include empirically estimated rates of error (Section S4.1); in the following referred to as data set $\mathcal{B}'$. A copy of this data set was further modified by performing *in silico* haplotype phasing (Section S4.2); in the following referred to as data set $\mathcal{B}''$. The process of generating data sets $\mathcal{B}$, $\mathcal{B}'$, and $\mathcal{B}''$ is summarized in Figure S17.

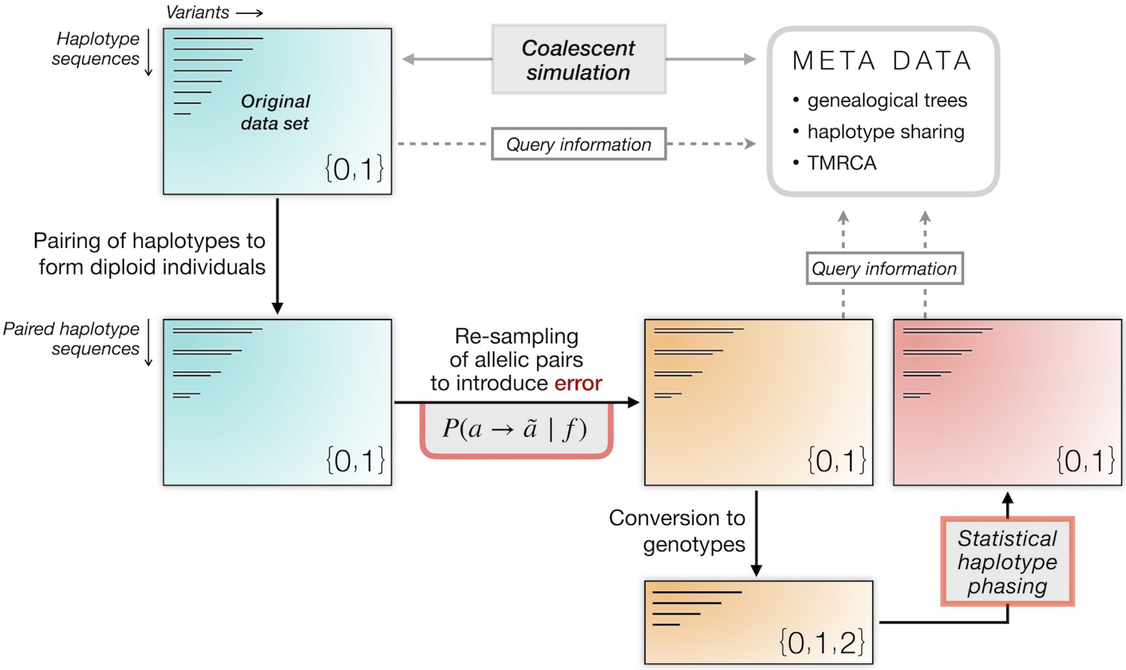

**Figure S17. Schematic of the data generation process.** The figure summarizes the generation of data sets $\mathcal{B}$ (*blue*), $\mathcal{B}'$ (*orange*), and $\mathcal{B}''$ (*red*). We performed coalescent simulations using `msprime` software [13], which produced a sample of haplotype sequences and a corresponding record (*meta data*) of the genealogical history of the simulated sample. Haplotype data was stored in *variant call format* (VCF), in which haploid sequences were arranged in pairs to form diploid individuals. We used this sample configuration to introduce error by scanning along the paired sequence in each individual, but where we used the frequency-dependent genotype error profile constructed in Section S3 to introduce error on haplotype-level. A given allelic pair, $a$, was re-sampled as $\tilde{a}$ with probability $P(a \to \tilde{a}|f)$, where $f$ is the sample allele frequency at the current site; see Equation (47). We further modified the resulting haplotype data set by performing *in silico* haplotype phasing using `SHAPEIT2` [5] after conversion to genotype data. Because these data were derived from the same original data set, its simulation record can be queried in downstream analyses of each generated data set. However, note that genealogical information may not be retrieved conclusively in data after haplotype phasing.

**Simple demographic model simulation ($\mathcal{A}$).** We simulated a sample of $N$=1,000 haplotypes of 100 Mb length, with $N_e$=10,000 and constant and equal mutation and recombination rates ($\mu = 1 \times 10^{-8}$, $r = 1 \times 10^{-8}$, per base per generation). The full command to simulate this data set is given in Script 1 (below).

**Script 1.** Simple demographic model; single demographic unit without migration, constant population size, and constant and equal mutation and recombination rates.

```
import msprime
data = msprime.simulate(Ne = 10000,
                        sample_size = 1000,
                        mutation_rate = 1e-08,
                        recombination_rate = 1e-08,
                        length = 100000000)
data.dump("history.hdf5") # simulation record
with open("sample.vcf", "w") as vcf_file:
        data.write_vcf(vcf_file, ploidy = 2) # output haplotype data in VCF file
```

**Complex demographic model simulation ($\mathcal{B}$).** We simulated a sample of $N = 5,000$ haplotypes ($N_e = 7,300$; $\mu = 2.35 \times 10^{-8}$ per base per generation) under a demographic model that recapitulates the human expansion out of Africa, following Gutenkunst *et al.* [1], with population growth and migration between three major populations (African, Asian, European). The simulation was conducted with variable rates of recombination, for which we used the genetic map for Chromosome 20 from HapMap (Phase 2; GRCh37) [2], which also determined the length of the simulated region (~63 Mb). The full command to simulate this data set is given in Script 2 (below).

**Script 2.** Complex demographic model; out-of-Africa model following [1], with population growth and migration between three major populations (African, Asian, European), with constant mutation rate, and variable recombination rates in Chromosome 20 from HapMap (Phase 2; GRCh37) [2]. Script modified from the msprime manual (https://msprime.readthedocs.io).

```
import msprime
import math
rec_map = msprime.RecombinationMap.read_hapmap("genetic_map_GRCh37_chr20.txt")
mut_rate = 2.35e-8
N_A = 7300
N_B = 2100
N_AF = 12300
N_EU0 = 1000
N_AS0 = 510
generation_time = 25
T_AF = 220e3 / generation_time
T_B = 140e3 / generation_time
T_EU_AS = 21.2e3 / generation_time
r_EU = 0.004
r_AS = 0.0055
N_EU = N_EU0 / math.exp(-r_EU * T_EU_AS)
N_AS = N_AS0 / math.exp(-r_AS * T_EU_AS)
m_AF_B = 25e-5
m_AF_EU = 3e-5
m_AF_AS = 1.9e-5
m_EU_AS = 9.6e-5
population_configurations = [
    msprime.PopulationConfiguration(sample_size=0, initial_size=N_AF),
    msprime.PopulationConfiguration(sample_size=5000, initial_size=N_EU, growth_rate=r_EU),
    msprime.PopulationConfiguration(sample_size=0, initial_size=N_AS, growth_rate=r_AS)
]
migration_matrix = [
    [      0, m_AF_EU, m_AF_AS],
    [m_AF_EU,       0, m_EU_AS],
    [m_AF_AS, m_EU_AS,       0],
]
demographic_events = [
    msprime.MassMigration(time=T_EU_AS, source=2, destination=1, proportion=1.0),
    msprime.MigrationRateChange(time=T_EU_AS, rate=0),
    msprime.MigrationRateChange(time=T_EU_AS, rate=m_AF_B, matrix_index=(0, 1)),
    msprime.MigrationRateChange(time=T_EU_AS, rate=m_AF_B, matrix_index=(1, 0)),
    msprime.PopulationParametersChange(time=T_EU_AS, initial_size=N_B, growth_rate=0, population_id=1),
    msprime.MassMigration(time=T_B, source=1, destination=0, proportion=1.0),
    msprime.PopulationParametersChange(time=T_AF, initial_size=N_A, population_id=0)
]
data = msprime.simulate(Ne = N_A,
                        mutation_rate = mut_rate,
                        recombination_map = rec_map
                        population_configurations = population_configurations,
                        migration_matrix = migration_matrix,
                        demographic_events = demographic_events)
data.dump("history.hdf5") # simulation record
with open("sample.vcf", "w") as vcf_file:
        data.write_vcf(vcf_file, ploidy = 2) # output haplotype data in VCF file
```

#### S4.1 Integration of error in simulated data

The frequency-dependent genotype error profile constructed in Section S3 was used to modify a copy of simulated data set $\mathcal{B}$, but where we introduced error on haplotype-level. This modified version is referred to as data set $\mathcal{B}'$. We first sorted haplotypes into pairs to form diploid individuals composed of two allelic sequences. Note that `msprime` already provides the option to output simulated haploid sequences in *variant call format* (VCF) for diploids. Alleles are encoded as 0s and 1s for the ancestral and derived allelic states, respectively. There are four possible ordered pairs of alleles; namely

$$a_{00} = (0,0),\ a_{01} = (0,1),\ a_{10} = (1,0),\ a_{11} = (1,1).$$

We scanned along the paired sequence of each individual, where, for a given allelic pair $a$, we sampled $\tilde{a}$ from the set of possible pairs $\{a_{ij}\}_{i,j\in\{0,1\}}$, with probability according to the empirically determined rate of genotype error; given by

$$P(a \to \tilde{a}|f) = \begin{array}{c} \begin{matrix} \tilde{a}_{00} & \tilde{a}_{01} & \tilde{a}_{10} & \tilde{a}_{11} \end{matrix} \\ \begin{bmatrix} e_{0\to0} & \frac{e_{0\to1}}{2} & \frac{e_{0\to1}}{2} & e_{0\to2} \\ e_{1\to0} & e_{1\to1} & 0 & e_{1\to2} \\ e_{1\to0} & 0 & e_{1\to1} & e_{1\to2} \\ e_{2\to0} & \frac{e_{2\to1}}{2} & \frac{e_{2\to1}}{2} & e_{2\to2} \end{bmatrix} \begin{matrix} a_{00} \\ a_{01} \\ a_{10} \\ a_{11} \end{matrix} \end{array} \tag{47}$$

where $f$ is the allele frequency in the simulated sample at the current site along the sequence. Recall that the coefficient $e_{i\to j}$ captures the relative rate of error per genotypic class at a set of recorded allele frequencies; see Equation (46). At sites where the observed frequency did not match to frequencies recorded in the error profile, we used linear interpolation to approximate error rates, which we again normalized to sum to 1 per true genotype class (corresponding to rows in Equation 47).

#### S4.2 Additional haplotype error through *in silico* phasing

We used a copy of data set $\mathcal{B}'$ (described above) to introduce additional errors through *in silico* haplotype phasing. Note that haplotype sequences were already sorted to form diploid individuals in the generated VCF output, and we maintained this configuration after the integration of data error. We computed the genotype sequence per individual as the sum of alleles at each position. This converted data set was then used to statistically re-estimate haplotypes without a reference panel using `SHAPEIT2` [5]. The haplotype phasing process was expected to introduce single-site ("flip") and long-range phase ("switch") errors.

### S5 Shared haplotype estimation using a simple hidden Markov model (HMM)

We have developed a hidden Markov model (HMM) to locally infer the region a given pair of chromosomes (concordant or discordant) share by descent from their MRCA at a focal position. Relative to a given target site in the genome, the shared haplotype segment is delimited by ancestral recombination events that occurred independently to the left and right-hand side in either of the two lineages considered. Our HMM is constructed as a two-state model in which the *local* genealogy at a given target site is distinguished from any *peripheral* genealogies that generated the variation seen outside the local segment. The two states are denoted by $H_0$ (local) and $H_1$ (peripheral). We scan the sequence from the position of a given target site until the end of the chromosome, which we do in two independent runs for the sequence to the left and the sequence to the right-hand side. The Viterbi algorithm is then used to decode the hidden state sequence from which we find a recombination breakpoint at the first occurrence of the $H_1$ state. If the $H_0$ state was inferred at all sites until the end of the chromosome, we record the breakpoint to sit beyond the last position of the sequence. The breakpoints inferred on both sides identify the sequence interval which encloses the underlying shared haplotype segment. This is illustrated in Figure S18. In cases where the $H_1$ state was inferred throughout (including the focal site at the initial position), we exclude the current (concordant or discordant) haplotype pair from downstream analyses.

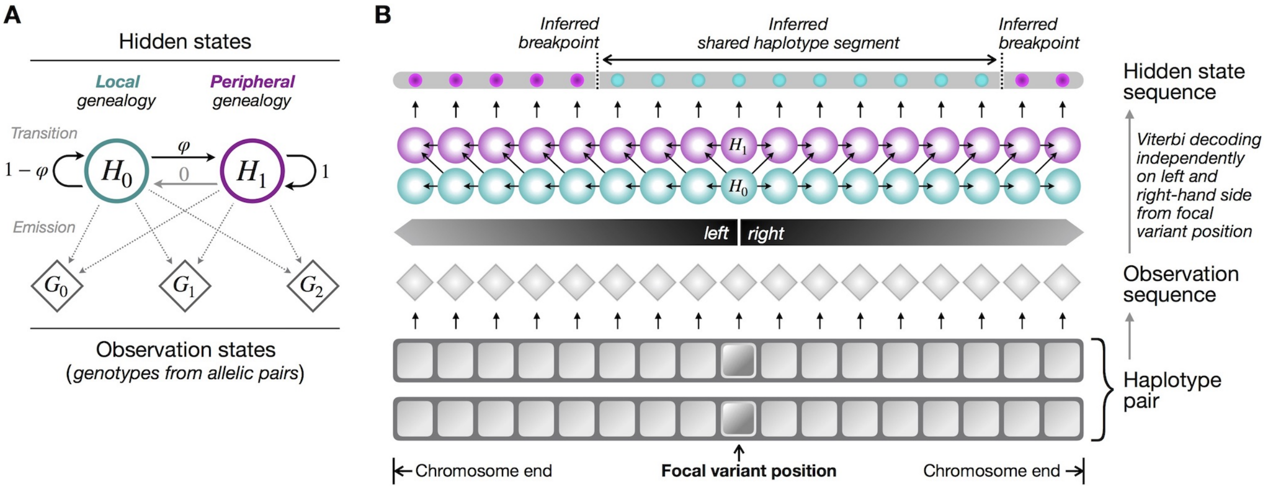

**Figure S18. Schematic of the hidden Markov model (HMM).** Panel **A** gives a graphic representation of the hidden state transitions and observation state emissions of the HMM. We define two hidden states to discriminate the *local* genealogy ($H_0$) which generated the variation observed at and around a given target site from any *peripheral* genealogies ($H_1$) outside the focal segment. The observation sequence is encoded as genotypes ($G_0, G_1, G_2$) by combining alleles (encoded as 0s and 1s) along the sequence of the haplotype pair considered. The probability of transition from the local to the peripheral state is denoted by $\varphi$. The HMM is constructed using a *left-to-right* architecture in which transitions from a higher to a lower-numbered state have zero probability. Emission probabilities were determined using an empirically generated model with realistic rates of data error. Panel **B** shows the HMM trellis, illustrating (*from bottom to top*) how two haplotype sequences are paired to form the observation sequence. Starting at the position of a given focal variant, the HMM is applied independently to the sequence to the left and right-hand side, to infer the nearest recombination *breakpoints* that delimit the enclosed shared haplotype segment.

The observation sequence is constructed from SNP variant data from the two sequences considered. Alleles are encoded as 0s and 1s, denoting the reference and alternate allele, respectively. We assume that the alternate allelic state has been correctly assigned to the derived allele. The two sequences are paired such that each site is represented as an unsorted pair of alleles (genotypes) with three possible observation states; namely

$$G_0 = \{0,0\},\ G_1 = \{0,1\},\ G_2 = \{1,1\}.$$

Note that we do not consider a separate state for missing alleles, but rather exclude sites from the observation sequence at which one or both alleles have been missed.

##### S5.1 Transition model

Our HMM employs a *left-to-right* architecture, meaning that the state transitions proceed in one direction where transitions from a higher to a lower-numbered state have zero probability. We define the transition matrix

$$A \;=\; \begin{bmatrix} a_{00} & a_{01} \\ a_{10} & a_{11} \end{bmatrix} \;=\; \begin{bmatrix} 1-\varphi & \varphi \\ 0 & 1 \end{bmatrix} \tag{48}$$

where the coefficient $a_{ij}$ denotes the probability of transition from state $H_i$ to state $H_j$. Since we have $a_{10} = 0$ (and thus $a_{11} = 1$), it is implied that we cannot return to the local genealogy once it has been left. The transition from the local to the peripheral state is given by $a_{01} = \varphi$, which is dependent on the genetic distance between consecutive variants in the sequence and the number of meioses (generations) separating the two haplotypes considered.

From the target site at position $k$, the HMM proceeds until the end of the chromosome to either the left or right-hand side relative to $k$. At the current site $l$ in the sequence, we define $\delta_l$ as the genetic distance observed between $l$ and the immediately previous site at position $l-1$. The time since the two chromosomes inherited the local haplotype from a common ancestor is treated as an unknown, but we use the following approximation to make broad distinctions of coalescence times based on the sample frequency $f_k$ of the allele observed at target site $k$. For concordant pairs, we calculate [14]

$$\xi_k \;=\; \frac{-2f_k}{1-f_k}\,\log(f_k)\,, \quad 0 < f_k < 1 \tag{49}$$

which is the expected age of a selectively neutral allele at frequency $f_k$ in a population of constant size. Since the above is dependent on the frequency of the allele observed at the target site, it does not apply to discordant pairs, for which we set $\xi_k = 1$, as we cannot derive an expectation when the allele is not shared by both haplotypes. We use the above to obtain an approximation for the probability of transition from the local to a peripheral genealogy as

$$\varphi_l(k) \;=\; 1 - e^{-4N_e\delta_l\xi_k} \tag{50}$$

where $N_e$ is the diploid effective size of the population. Note that we therefore compute a transition matrix $A_l(k)$ at every site along the sequence, dependent on the allele frequency at the target site $k$ and the genetic distance at the current site $l$.

Transition probabilities computed for different focal allele frequencies are illustrated in Figure S19A. We note that differences in transition probabilities resulting from focal allele frequency variation, in practice, have little impact on the allele age estimation process. This is demonstrated through analysis of simulated data set $\mathcal{A}$ (see Section S4). First, we calculated transition probabilities conditional on focal allele frequency, as defined in this section and which is the default in GEVA. We then analyzed the same data, but set $\xi_k = 1$ for all haplotype pairs to make the calculation of transition probabilities independent of frequency. We found that age estimates were highly correlated (Spearman rank correlation, $\rho > 0.98$) for each clock model; see Figure S19B.

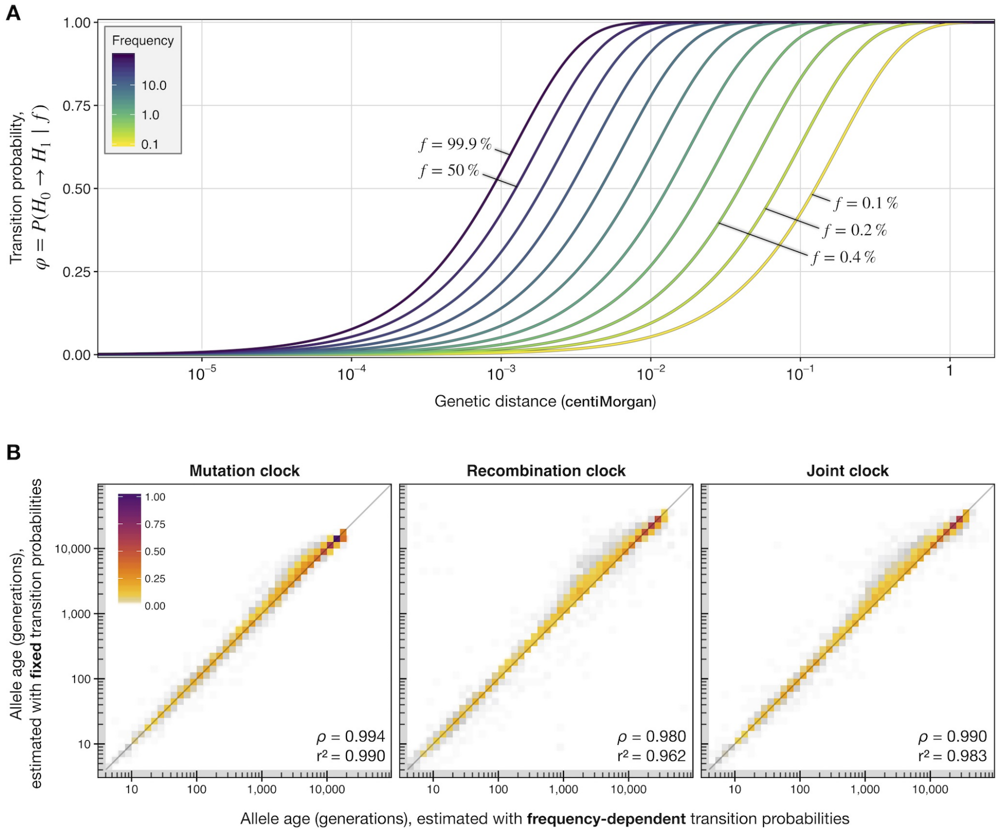

**Figure S19. Transition probabilities.** Panel **A** shows the probability of transition from the local ($H_0$) to the peripheral ($H_1$) state, $\varphi$, given the genetic distance (*x-axis*) for different allele frequencies (see legend; labels indicate certain low and high frequencies). Note that the genetic distance is given in *centiMorgan* in the figure, but transition probabilities are computed in units of *Morgan* in Equation (50). In the HMM, genetic distance is measured between consecutive sites along the sequence, and allele frequency is taken as the observed frequency of the alternative allele in the sample at a given focal site. Panel **B** shows density scatterplots of allele age estimated ($\max_{\mathcal{C}} = 500$; $\max_{\mathcal{D}} = 500$) under each clock model for 5,000 variants (randomly drawn at allele count $1 < x < N$) from data simulated with sample size $N$=1,000 haplotypes, $N_e$=10,000, $\mu = 1 \times 10^{-8}$, and $r = 1 \times 10^{-8}$; see Section S4 (Script 1). The same set of variants was analyzed using GEVA, but with frequency-dependent transition probabilities in the HMM (*x-axis*), and with fixed transition probabilities (*y-axis*). For the latter, we set $\xi_k = 1$ for all haplotype pairs; see Equation (50). Lower inserts indicate the Spearman rank correlation statistic, $\rho$, and the squared Pearson correlation coefficient (on log-scale), $r^2$, calculated between the two corresponding sets of age estimates.

#### S5.2 Empirical emission model

We generated an empirical model from simulated data, which we modified to include realistic distributions of error, to make the HMM robust in applications to real data. The model is defined by a set of observation probabilities (emissions), denoted by $\sigma_j^i(f_l)$, where $i$ identifies the hidden state $\{H_i\}_{i\in\{0,1\}}$ and $j$ the genotypic state $\{G_j\}_{j\in\{0,1,2\}}$ observed at site $l$ along the sequence of the two haplotypes considered. Emissions are obtained conditional on the allele frequency, $f_l$, observed in the sample at site $l$. Note that the emissions for the two hidden states ($H_0$, $H_1$) are based on TMRCA, but which involves a simplifying assumption (described further below) to construct a time-independent, approximate emission model.

The model was constructed by estimating the relative rates of observing each genotypic state in pairwise shared haplotype segments (identified from genealogical records after simulation using `msprime`; see Section S4), given allele frequency information and the TMRCA for each segment. This was done through an iterative sampling process. We randomly selected two haplotypes from simulated data and identified the locations of recombination breakpoints to detect shared haplotype segments (non-recombinant sequence intervals). Specifically, we scanned each genealogical tree along the sequence and recorded breakpoints at coordinates where the genealogical relationship changed due to recombination, such that the sequence interval in between two consecutive breakpoints was derived from the same MRCA. Note that the continuous position coordinates internally stored by `msprime` may differ from the discrete genomic positions after output to variant call format (VCF). We matched breakpoint coordinates to the returned genomic positions to identify sequence intervals that are observable from the data. For each segment, we recorded the TMRCA and retrieved the enclosed variant sequences from both haplotypes. The genotypic states observed at each site in the sequence interval were recorded together with the sample allele frequency at each site. Segments shorter than 10 variant sites were removed. We repeated this process until collecting $\sim$100 million segments (from >30,000 random haplotype pairs).

To illustrate differences arising from data error, we first analyzed data set $\mathcal{B}$, which was simulated using `msprime` under a complex demographic model that recapitulates the human expansion out of Africa (described in Section S4). This is compared to data set $\mathcal{B}'$, which is a copy of $\mathcal{B}$ but where haplotypes were modified using empirically estimated error rates (described in Section S4.1). Both data sets thereby have the same genealogical history.

We pooled segments into 100 TMRCA bins (evenly distributed on log-scale between 1 generation and 500,000 generations), within which we pooled genotypes into 500 allele frequency bins (evenly distributed on linear scale). For each combination of TMRCA and frequency bin, the relative rate of observing each genotypic state was obtained by normalizing counts (to sum to 1). The resulting time and frequency-dependent distributions are shown in Figure S20A. We found that the main differences were located at the extremes of either TMRCA or frequency. But notably, before error, the relative rate of observing heterozygous genotypes ($G_1$) was zero (or near zero) in each frequency bin for recent TMRCA (<10 generations), but non-zero throughout after error.

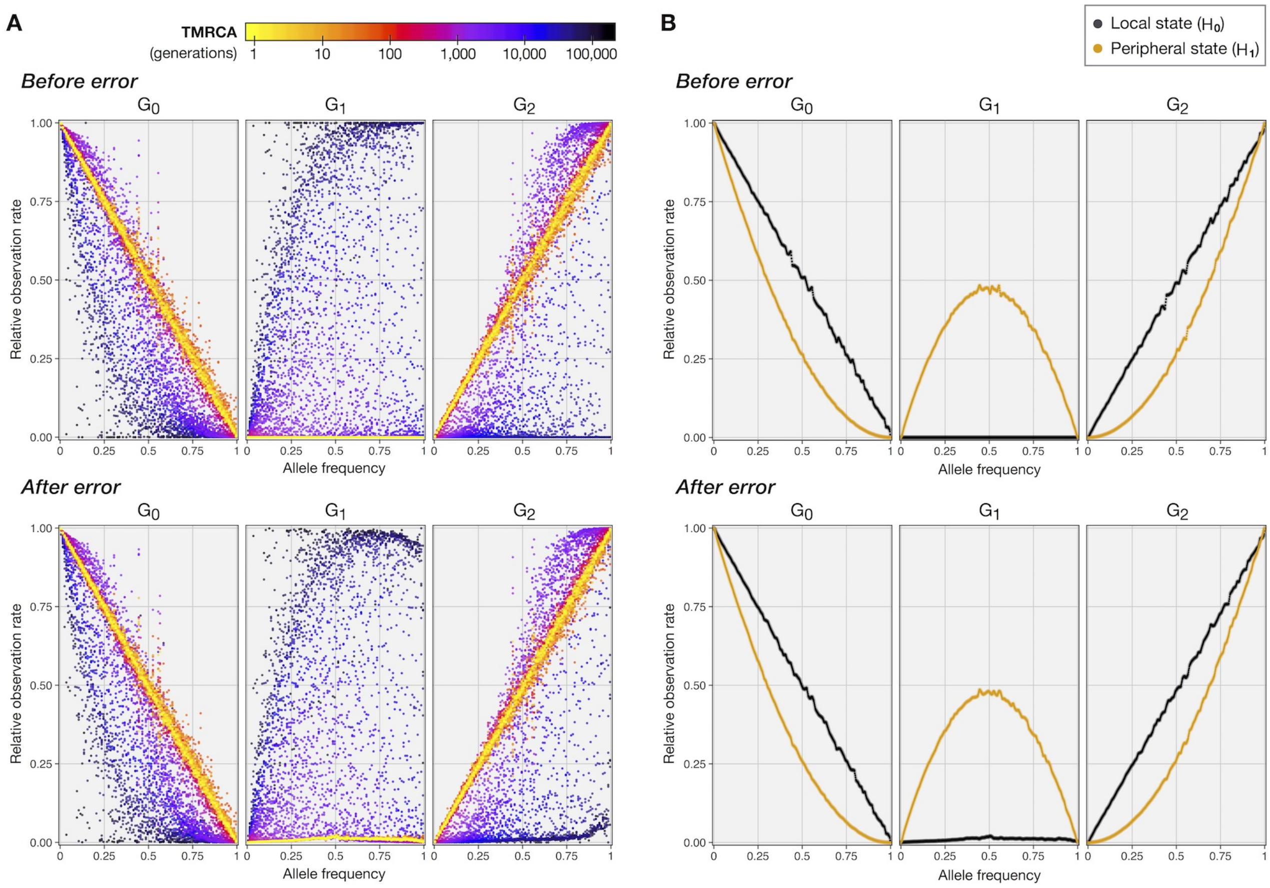

**Figure S20. Empirical emission probabilities.** The relative rate of observing each genotypic state $(G_0, G_1, G_2)$ in simulated data before error (data set $\mathcal{B}$; *top*) and after error (data set $\mathcal{B}'$; *bottom*). Panel **A** shows the relative rates measured in 100 TMRCA bins, evenly distributed on log-scale between 1 generation and 500,000 generations (see legend), and for variants pooled into 500 evenly distributed allele frequency bins. For each combination of TMRCA and frequency bin, measured rates were normalized to sum to 1 across the three genotypic states. Panel **B** shows the relative rates as in (A), but where a nominal distinction at 100 generations was made to determine emission probabilities in the local state ($H_0$; $\leq 100$ generations) and the peripheral state ($H_1$; $> 100$ generations) by allele frequency.

Next, we estimated emissions as defined for the model, specifically $\sigma^i_j(f_l)$, for which we pooled segments into only two TMRCA bins, to make an approximate distinction between recent shared ancestry and older genealogical relationships. We applied a nominal cutoff at 100 generations to distinguish segments in hidden state $H_0$ (TMRCA $\leq 100$ generations) from those in hidden state $H_1$ (TMRCA $> 100$ generations). The resulting observation rates (after pooling into 500 allele frequency bins) are shown in Figure S20B, for the analyses before and after error. We used the observation rates obtained from data after error as the emission model in subsequent applications of GEVA. For frequencies not captured by the set of recorded frequency bins, in practice, we use linear interpolation to approximate the emissions at the observed sample allele frequency, which we again normalize to ensure that observation rates sum to 1 per hidden state.

Note that the distinction of TMRCA to determine observation rates in $H_0$ and $H_1$ is arbitrary for any nominal cutoff. We see this as a useful simplification to distinguish a *local* shared haplotype segment from the background “noise” of variation produced through *peripheral* genealogies. However, one caveat is that we implicitly assume that local TMRCA is on average younger than TMRCA at neighboring segments. Inference of breakpoints is expected to be less problematic at older segments if the local segment sits within an extended region where variation is derived from similarly old relationships, because we would quickly transition into the peripheral state and infer a breakpoint nearby. But, conversely, it is less likely that we can accurately infer breakpoints that delimit an older and short local segment if the immediately following segment is younger and relatively long.

Further, we note that a nominal cutoff at 100 generations, in practice, may not reflect a limit on the ability to infer breakpoints at shared haplotype segments with higher or lower TMRCA, or on the performance of the method to estimate allele age. This is demonstrated through comparison of age estimation in GEVA, performing pairwise analyses of the HMM with emission models generated from different nominal cutoffs of TMRCA. We used the default emission model generated with a TMRCA cutoff at 100 generations (from data after error) and prepared a second model in the same way, but with a cutoff at 1,000 generations. For the same set of 5,000 randomly selected variants in data set $\mathcal{B}'$, we found that age estimates were highly correlated (Spearman rank correlation, $\rho > 0.94$) for each clock model; see Figure S21.

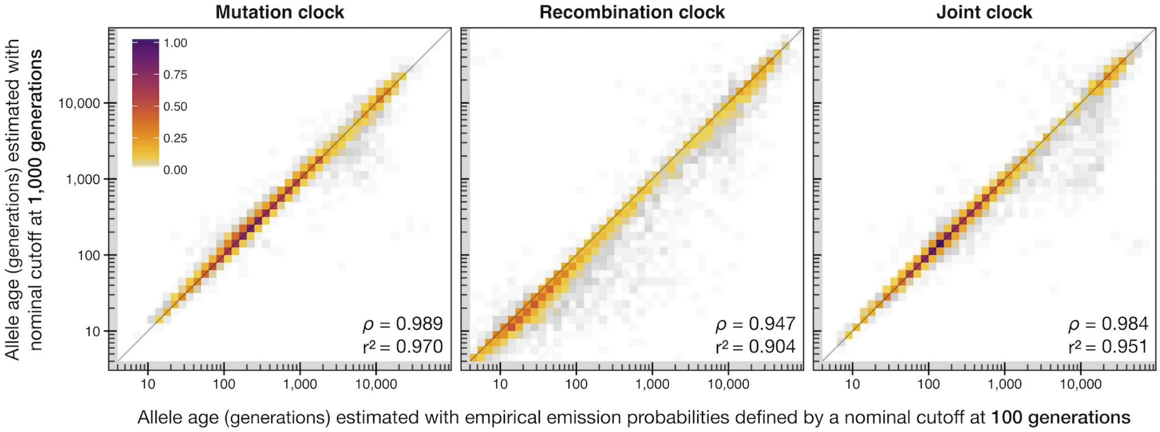

**Figure S21. Comparison of allele age estimated with different empirically generated emission models.** The correlation between allele age estimated using GEVA with two different emission models in the HMM. Emissions were generated from simulated data (after including realistic distributions of error), but with different nominal cutoffs to distinguish the two hidden states ($H_0$ and $H_1$) based on younger and older TMRCA, respectively. This was done by applying a TMRCA cutoff at 100 generations (*x-axis*) and with a cutoff at 1,000 generations (*y-axis*). Each panel shows the density scatterplots of allele age estimated ($\max_{\mathcal{C}} = 500$; $\max_{\mathcal{D}} = 500$) under a given clock model, for the same set of 5,000 variants (randomly drawn at allele count $1 < x < N$) from data set $\mathcal{B}'$, simulated with sample size $N$=5,000 haplotypes, $N_e$=7,300, $\mu = 2.35 \times 10^{-8}$, and variable recombination rates (HapMap2, Chromosome 20); see Section S4 (Script 2). Lower inserts indicate the Spearman rank correlation statistic, $\rho$, and the squared Pearson correlation coefficient (on log-scale), $r^2$, calculated between the two corresponding sets of age estimates.

#### S5.3 Empirical initial state model

Similar to the generation of emission probabilities described above, we estimated the initial probability of being in either $H_0$ or $H_1$ empirically from simulated data. For concordant pairs, we define $\pi_i^{\mathcal{C}}(f_k)$ as the initial probability of being in state $\{H_i\}_{i\in\{0,1\}}$, where $f_k$ is the sample allele frequency at the focal site $k$. Likewise, we define $\pi_i^{\mathcal{D}}(f_k)$ for discordant pairs. Again, we used simulation data sets $\mathcal{B}$ and $\mathcal{B}'$ (described in Section S4 and Section S4.1, respectively) to obtain frequency-dependent estimates for $\pi_i^{\mathcal{C}}$ and $\pi_i^{\mathcal{D}}$ from the true positive rate (TPR) of correctly observing allelic combinations by comparing pairs ($\{1,1\}$ for concordant pairs, $\{0,1\}$ or $\{1,0\}$ for discordant pairs) before and after error.

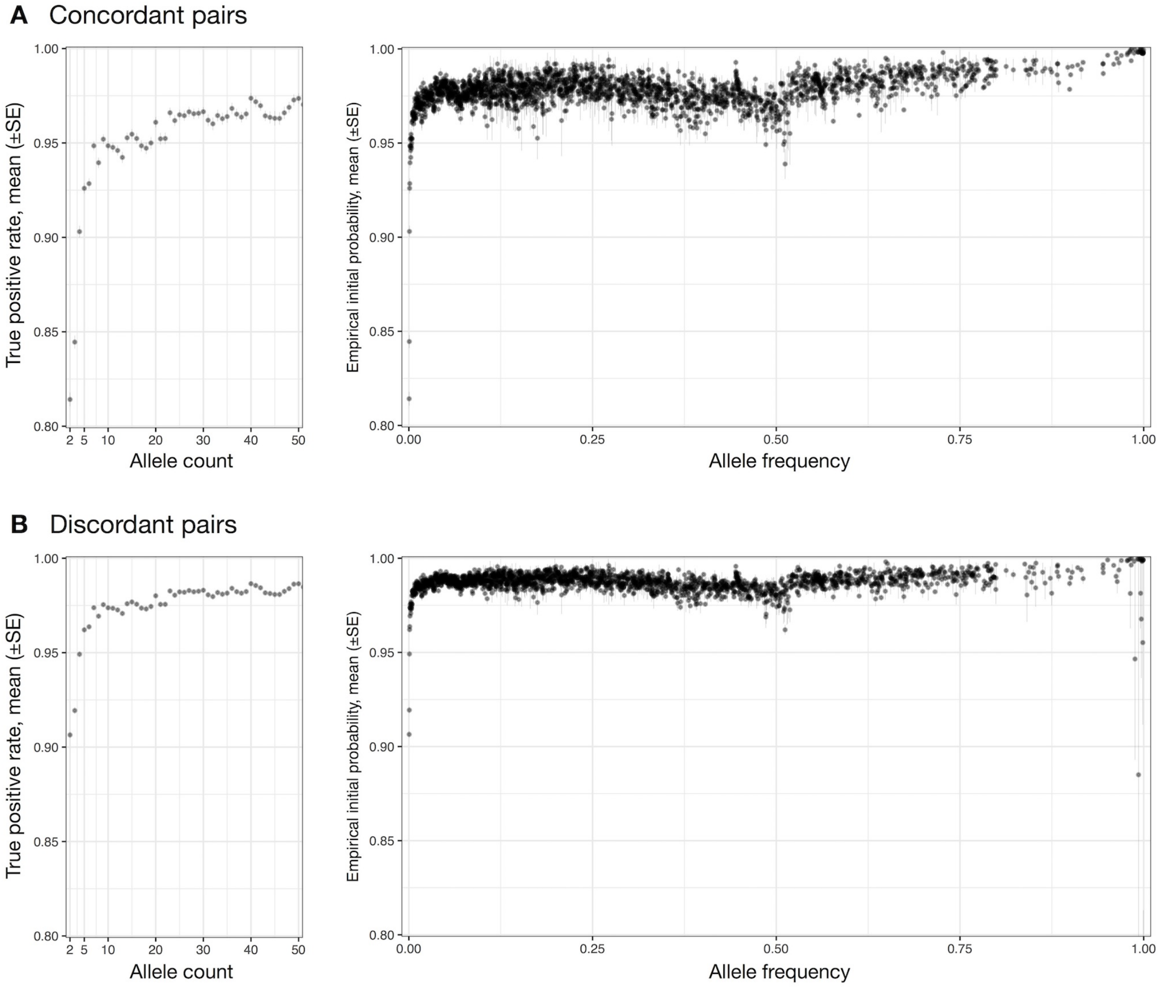

**Figure S22. Empirical initial state probabilities.** The probability of being in the local $H_0$ state at the initial position in the sequence was estimated from the true positive rate (TPR) of correctly observing the allelic combination $\{1,1\}$ for concordant and $\{0,1\}$ or $\{1,0\}$ for discordant pairs, by comparing the same pairs in data before error (data set $\mathcal{B}$) to corresponding coordinates in data after error ($\mathcal{B}'$). The TPR was measured at sets of up to 1,000 variants randomly selected per allele count category (1 $<$ allele count $< N$; where $N$=5,000 is the number of haplotypes in the simulated sample), for which up to 1,000 concordant or discordant pairs were sampled per variant. Panel **A** shows the mean TPR (±SE) per allele count category measured for concordant pairs; shown in greater detail for allele counts in $[2, 50]$ (*left*), and across the full allele frequency range (*right*). Panel **B** shows the same as **A**, but for discordant pairs.

In data without error, we have $\pi_0^{\mathcal{C}} = 1$ and $\pi_0^{\mathcal{D}} = 1$ with certainty at any $f_k$ (and therefore $\pi_1^{\mathcal{C}} = 0$ and $\pi_1^{\mathcal{D}} = 0$). To estimate initial state probabilities in data after error, we used data set $\mathcal{B}'$ to divide the set of variants by allele count (simulated sample size $N$=5,000). We then randomly sampled up to 1,000 sites per allele count category ($1 <$ allele count $< N$) and, for each site, sampled up to 1,000 pairs per set of possible concordant and discordant pairs. For both groups, we computed the TPR by comparison to the allelic combination seen at the same position and for the same pair in data before error (data set $\mathcal{B}$). We recorded the TPR at each site and report mean TPR per allele count category; see Figure S22.

We defined $\pi_0^{\mathcal{C}}(f_k)$ and $\pi_0^{\mathcal{D}}(f_k)$ according to the recorded mean TPR at a given allele frequency, and set $\pi_1^{\mathcal{C}}(f_k) = 1 - \pi_0^{\mathcal{C}}(f_k)$ and $\pi_1^{\mathcal{D}}(f_k) = 1 - \pi_0^{\mathcal{D}}(f_k)$. Because these were recorded at a fixed set of allele frequencies, in practice, we use linear interpolation to approximate the probabilities at sample allele frequency $f_k$, which we again normalize to ensure that probabilities sum to 1 per hidden state, separately for concordant and discordant pairs.

### S6 Simulation study

We used simulated data to assess the genealogical estimation of variant age (GEVA) method (Section S6.1), and compared GEVA to a strategy that employs PSMC as a method to infer TMRCA posteriors from which allele age can be estimated (Section S6.1.1). Also, we evaluated the performance of the heuristic rejection method through which outliers in the inferred TMRCA distribution are excluded prior to age estimation in GEVA (Section S6.1.2). Lastly, we assessed our method to infer the cumulative coalescent function (CCF) by comparison to the true ancestry shared among samples in simulated data (Section S6.2).

#### S6.1 Estimation of variant age in simulated sample data

The performance of GEVA was evaluated in terms of its accuracy to infer the TMRCA between pairs of haplotypes, and to estimate allele age from the inferred TMRCA distributions. We used simulation data sets $\mathcal{A}$ and $\mathcal{B}$, which we generated under different demographic models (described in Section S4). To reproduce the conditions typically present in applications to real data, we also applied GEVA to data sets $\mathcal{B}'$ and $\mathcal{B}''$, which we derived from $\mathcal{B}$, but where haplotype data were modified to contain realistic proportions of data error ($\mathcal{B}'$; described in Section S4.1) and additional error through *in silico* haplotype phasing ($\mathcal{B}''$; described in Section S4.2).

We applied GEVA with scaling parameters as specified for each simulation. For data set $\mathcal{A}$, we used $N_e$ =10,000, $\mu = 1 \times 10^{-8}$, and $r = 1 \times 10^{-8}$. Allele age was estimated for 5,000 variants randomly drawn from sites with allele count $1 < x < N$, which we analyzed with $\max_{\mathcal{C}} = 500$ and $\max_{\mathcal{D}} = 500$ as the maximum number of concordant and discordant pairs sampled per site (given a sample size of $N$ =1,000). Thus, we inferred TMRCA posteriors from locally estimated shared haplotype segments at >3 million concordant and discordant pairs. We used our heuristic rejection method (Section S1.5) to exclude outlier pairs before estimating age.

Similarly, for $\mathcal{B}$ and its derived data sets $\mathcal{B}'$ and $\mathcal{B}''$, we used $N_e$ =7,300, $\mu = 2.35 \times 10^{-8}$, and the HapMap genetic map (Phase 2; GRCh37; Chromosome 20) to determine variable recombination rates over the simulated region. Data were simulated with a sample size of $N$ =5,000. Allele age was estimated for the same set of variants in $\mathcal{B}$, $\mathcal{B}'$, and $\mathcal{B}''$, with $\max_{\mathcal{C}} = 500$ and $\max_{\mathcal{D}} = 500$. We selected 5,000 variants at random from the intersection of sites at which the observed allele count satisfied $1 < x < N$ across the three data sets. More than 3 million haplotype pairs were analyzed in each data set, where we excluded outliers before estimating age.

The results of the analysis of data set $\mathcal{A}$ are shown in Figure S1, in which we used different metrics to measure estimation bias and correlation between true and estimated allele age (Figure S1A) and pairwise TMRCAs inferred at each site (Figure S1B), for each of the three clock models. Likewise, the results for data set $\mathcal{B}$ are shown in Figure S2, data set $\mathcal{B}'$ in Figure S3, and data set $\mathcal{B}''$ in Figure S4. The metrics are described below.

**Evaluation of allele age.** Let $t$ denote the true time of the mutation event that gave rise to the allele at a given site in the sample, for which we obtain an estimated age denoted by $\hat{t}$. Coalescent simulators typically do not record the actual point in time when a (neutral) mutation occurred, as the probability of mutation is proportional to the branch length in a simulated genealogical tree, and mutations are placed uniformly over a given branch. To obtain a "true" value for allele age, we queried the simulation meta-data as recorded by `msprime` to locate the branch on which a specific mutation occurred. A branch is delimited by the time all carrier haplotypes have coalesced into a single lineage and the time this lineage joined with any of the remaining lineages. Let $t^0$ and $t^1$ denote the times of coalescent events that occurred immediately before and after the mutation event, respectively, as for the simulated sample, where $t^0 \leq t$ and $t^1 > t$. We retrieved $t^0$ and $t^1$ from simulation records to calculate the true time of a mutation as $t' = \sqrt{t^0 \times t^1}$, which is the geometric mean. Note that the arithmetic mean would be appropriate given that mutation events are placed uniformly between $t^0$ and $t^1$ in the neutral coalescent. However, we found the geometric mean to be more reliable when analyzing larger sets of variants that exhibit high variability in terms of mutational timing and branch lengths.

Correlation between true ($t'$) and estimated ($\hat{t}$) allele age was measured using Spearman's rank correlation coefficient, $\rho$. Additionally, we calculated Pearson's $r^2$ on log-scaled values of true and estimated allele age. Estimation error was measured using the root mean squared $\log_{10}$ error (RMSLE), calculated as

$$\text{RMSLE} = \sqrt{\frac{1}{n} \sum_{i=1}^{n} \Big( \log_{10} \left[\, t'_i \,\right] - \log_{10} \left[\, \hat{t}_i \,\right] \Big)^2} \tag{51}$$

where $n$ is the number of variants considered.

We compute the following metric as a measure for estimation bias, denoted by $\epsilon$, which is adjusted for the time interval between $t^0$ and $t^1$. Error is quantified as the mean difference relative to the time interval during which a focal mutation occurred, where underestimation of allele age is calculated relative to $t^0$, and overestimation relative to $t^1$; calculated as

$$\epsilon = \frac{1}{n} \sum_{i=1}^{n} I_i^0 \left( \frac{t_i^0 - \hat{t}_i}{t_i^0} \right) + I_i^1 \left( \frac{\hat{t}_i - t_i^1}{t_i^1} \right) \tag{52}$$

where $I^0$ and $I^1$ are indicator functions given by

$$I_i^0 = \begin{cases} 0 \ \text{ if } \ t_i^0 < \hat{t}_i \\ 1 \ \text{ otherwise} \end{cases} , \ I_i^1 = \begin{cases} 0 \ \text{ if } \ t_i^1 > \hat{t}_i \\ 1 \ \text{ otherwise} \end{cases} . \tag{53}$$

Implicitly, this considers any age estimate falling in between $t^0$ and $t^1$ as being correct, thereby making $\epsilon$ robust towards variations in branch lengths when considering larger sets of variants with different coalescent histories.

**Evaluation of pairwise TMRCA.** We used Spearman's $\rho$ and Pearson's $r^2$ to measure correlation between true and inferred TMRCA for the sets of concordant and discordant pairs sampled for each variant, as well as RMSLE as an error metric. The true time of coalescence was determined from simulation records, and we used the mean of the Gamma distribution as a point estimate of the inferred TMRCA, calculated as $\alpha/\beta$, where the values of $\alpha, \beta$ were obtained through analysis under a given clock model (see Section S1.3).

###### S6.1.1 Variant age estimation based on PSMC

The *Pairwise Sequentially Markovian Coalescent* (PSMC) model has been used to infer historic changes in population size back in time [15], using sequence data from two haplotypes alone (or one diploid individual). The model is based on the *Sequentially Markov Coalescent* (SMC) model for analytically tractable approximation to the ancestral recombination graph (ARG) in model-based inferences [16, 17]. Here, we employed the HMM-based PSMC method [15] for TMRCA inference between two chromosomal sequences. In this approach, time is divided into a number of discrete intervals, which are the hidden states of the HMM. Using the forward-backward algorithm, a posterior probability is obtained for each state at different sites equally spread across the full length of the chromosome.

**Implementation.** Here we used an implementation of the PSMC-based HMM to infer the posterior distribution of coalescence times for concordant and discordant pairs at a given target position; thus effectively treating PSMC as a substitute clock model. The `decode` algorithm implemented in software available for the *Multiple Sequentially Markovian Coalescent* (MSMC) method [18] specifically applies the PSMC-HMM when two haplotype sequences are provided as input data. We modified `decode` to only output posterior probabilities at a specified target position, but without hindering the computation of posteriors along the sequence. The modified version of `decode` is available online.*

**Pairwise TMRCA inference.** We first used GEVA to estimate allele age for the sets of variants selected from each simulated data set ($\mathcal{A}, \mathcal{B}, \mathcal{B}', \mathcal{B}''$), to then apply the modified `decode` algorithm on haplotype data for the same sets of concordant and discordant pairs as sampled through GEVA (before excluding outlier pairs). The number of discrete time intervals (hidden states) was set to 64, which is the default in `decode`. For simulation panel $\mathcal{A}$, we used $N_e$=10,000, $\mu = 1 \times 10^{-8}$, and $r = 1 \times 10^{-8}$ to specify the scaled mutation and recombination rates. For $\mathcal{B}$, $\mathcal{B}'$, and $\mathcal{B}''$, we used $N_e$ =7,300 and $\mu = 2.35 \times 10^{-8}$ to specify the scaled mutation rate ($\theta = 2N_e\mu$). Because `decode` does not operate on variable recombination rates, we fixed the scaled recombination rate parameter to 80% of the value of $\theta$, which is the recommended setting.† A total of >14 million haplotype pairs was analyzed across all four simulated data sets.

* https://github.com/pkalbers/msmc2
† https://github.com/stschiff/msmc/blob/master/guide.md

**Allele age estimation.** We first applied our heuristic method to reject outlier pairs in the sets of concordant and discordant pairs that were analyzed per focal variant; see Section S1.5. For this, we took the mode of the inferred posterior distribution to obtain a point estimate of the TMRCA per pair, which we recorded at the mean between time interval boundaries. We then estimated allele age on the posteriors of the retained pairs using a composite posterior approach similar to the one used by GEVA (Section S1.4). Specifically, we computed

$$F(t) = \sum_{i=1}^{t} p(i) \times \max\left[\sum_{j=1}^{n} p(j)\right]^{-1}, \quad \text{for } t = 1, 2, \ldots, n \tag{54}$$

for concordant pairs and $F'(t_i) = 1 - F(t_i)$ for discordant pairs to approximate the cumulative posterior distribution, where $n$ is the number of discrete time intervals and $p(i)$ the posterior probability inferred at the $i$th time interval. The PSMC-derived composite posterior distribution was then computed as

$$\Phi(t) \;\propto \prod_{\{a,b\}\in C} F_{a,b}(t) \;\times \prod_{\{c,d\}\in D} F'_{c,d}(t)\;, \quad \text{for } t = 1, 2, \ldots, n \tag{55}$$

where $C$ and $D$ refer to the pre-selected (and subsequently filtered) sets of available concordant and discordant pairs, respectively. A point estimate of allele age was taken at the mode of the composite posterior distribution which we recorded at the mean between time interval boundaries.

##### S6.1.2 Performance of the heuristic pair rejection method

We evaluated our heuristic rejection method (Section S1.5) in terms of the proportion of haplotype pairs rejected in data before and after error, for each clock model and the PSMC-based approach. In data before error ($\mathcal{B}$), pairwise TMRCA inference under the joint clock model led to a rejection of 2.4% of pairs, across the 5,000 variants analyzed, which was similar for the mutation clock (2.3%) and higher for the recombination clock (3.4%). In data after error ($\mathcal{B}'$), 8.1% of pairs were rejected for the joint clock, 7.6% for the mutation clock, and 10.7% for the recombination clock. Additional phasing of data after error ($\mathcal{B}''$) showed no notable differences compared to the proportion of pairs rejected in data set $\mathcal{B}'$. When using PSMC to infer TMRCA, only 1.7% of pairs were rejected in the analysis on data before error ($\mathcal{B}$), but 8.9% in data after error ($\mathcal{B}'$ or $\mathcal{B}''$).

We further evaluated our heuristic filtering approach in terms of its accuracy to reject pairs that have been selected by GEVA due to data error. This was done by comparing the pairs selected in the analysis of data set $\mathcal{B}'$ to their true allelic configuration in haplotype data from $\mathcal{B}$; results are given in Table S4. For the 5,000 variants analyzed, we measured the true positive rate (TPR) of correctly retaining error-free pairs, which was >94% for concordant or discordant pairs and under each clock model. The true negative rate (TNR) of correctly rejecting erroneous pairs was overall lower and differed between the two groups; >58%

for concordant pairs and >38% for discordant pairs under each clock model. Filtering of pairs inferred using the PSMC-based approach resulted in a similar TPR overall (>94% for concordant or discordant pairs), but the TNR was much lower compared to either clock model (9% for concordant pairs and 12.2% for discordant pairs), such that PSMC (within the implementation used here) showed the lowest accuracy.

**Table S4. Accuracy of rejecting haplotype pairs in data after error.** The performance of our heuristic rejection method (Section S1.5) was evaluated in terms of its accuracy to reject erroneous haplotype pairs that have been selected by GEVA from data after error ($\mathcal{B}'$). Selected concordant and discordant pairs were distinguished into true and false positives by scanning their allelic configurations in data before error ($\mathcal{B}$). We report the true positive rate (TPR, or sensitivity) of correctly retaining error-free pairs, the true negative rate (TNR, or specificity) of correctly rejecting erroneous pairs, as well as the accuracy (ACC) as the sum of true positives and true negatives divided by the sum of all pairs; separately for the sets of concordant and discordant pairs selected across 5,000 variants analyzed under each clock model and the PSMC-based approach. However, because the 5,000 variants analyzed here were selected from the intersection of sites at which the observed allele count satisfied $1 < x < N$ across data sets $\mathcal{B}$, $\mathcal{B}'$, and $\mathcal{B}''$, the accuracy reported is likely to be artificially inflated overall.

| Clock model | Concordant pairs | | | Discordant pairs | | |
|---|---|---|---|---|---|---|
| | TPR | TNR | ACC | TPR | TNR | ACC |
| *Mutation clock* | 97.0% | 58.9% | 96.0% | 97.3% | 38.2% | 87.2% |
| *Recombination clock* | 94.1% | 66.6% | 93.3% | 94.3% | 41.6% | 85.3% |
| *Joint clock* | 97.3% | 60.0% | 96.3% | 96.8% | 39.7% | 87.0% |
| *PSMC-based approach* | 94.6% | 9.0% | 37.3% | 94.5% | 12.2% | 49.0% |

#### S6.2 Inference of shared ancestry in simulated sample data

We used `msprime` software [13] to simulate the demographic model illustrated in Figure S8A, which recapitulates the human expansion out of Africa [1]. This model was used previously in the generation of data set $\mathcal{B}$, but we now modified Script 2 (see Section S4, Page 37) to simulate a sample of $N = 600$ haplotypes, consisting of an equal number of 200 haplotypes from each of the three simulated populations; African (AF), Asian (AS), and European (EU). Other simulation parameters were left unchanged. The model specifies three relevant events in the past; the time of the split of EU and AS from ancestral population B ($T_{EU-AS} = 848$ generations ago), the emergence of B from AF ($T_B$ = 5,600 generation ago), and the emergence of AF from ancestral population A ($T_{AF}$ = 8,800 generation ago). Only AS and EU experienced exponential population growth following $T_{EU-AS}$.

The simulated region encompassed 622,240 variant sites. We used GEVA to estimate the age of 412,348 variants (all SNPs with allele count $1 < x < N$) with $\max_{\mathcal{C}} = 100$ and $\max_{\mathcal{D}} = 100$ as the maximum number of concordant and discordant pairs sampled per site, and with scaling parameters as specified for the simulation ($N_e$ =7,300; $\mu = 2.35 \times 10^{-8}$; variable recombination rates from HapMap Chromosome 20). Variant age estimated under the join clock model was used for the inference of shared ancestry. As for previous analyses of GEVA on simulated data, we first assessed the correlation between true and estimated allele age, which we found to be consistent with previous results; see Figure S8B.

We inferred the ancestry of the simulated sample between every pair of haplotypes. This was done in three ways:

- **True CCF.** Coalescent profiles were obtained directly from simulation records, where we scanned the ancestry of a pair of lineages to determine the exact lengths of chromosomal segments that have coalesced up to a given point back in time. The fraction of the genome shared was computed as the sum of segment lengths divided by the full length of the simulated region, where we obtained the cumulative distribution at the exact time points of coalescent events.
- **Inferred CCF from true allele age.** The CCF between target and comparator genomes was inferred using the dynamic programming method described in Section S2.1, but where we used the true age of alleles to obtain the age-sorted sequence of observed allele sharing. True age was determined from simulation records, which we computed as the geometric mean between the times of coalescent events that delimit the branch on which a given mutation occurred (see Page 48). We only considered those variants for which also the estimated age was available.
- **Inferred CCF from estimated allele age.** Coalescent profiles were inferred using the dynamic programming method with estimated allele ages (joint clock).

Inference of the CCF was done for every haplotype as target genome in turn against every haplotype in the sample (excluding itself) as comparator genome. True and inferred ancestry results were subsequently compared after approximating CCFs over a fixed grid of 500 time points between 1 and 500,000 generations ago (equally spaced on log-scale). Note that the dynamic programming method considers mutations carried by only the target genome. The inferred ancestry of target $i$ shared with comparator $j$ may therefore differ from the inferred ancestry of target $j$ with comparator $i$. This is not the case for true CCFs, because these were generated having full knowledge of the sample ancestry, such that the ancestry measured between target and comparator $i, j$ is identical to $j, i$.

The CCFs obtained for the three strategies are compared in Figure S8C, which shows the average fraction of the genome shared back in time for each combination of target and comparator population. The times of the simulated demographic events are reflected by changes of the gradient along the ancestry shared within and between different populations.

In results of the true CCF, the ancestry shared among only AS genomes increases rapidly (exponentially) back in time until reaching $T_{EU-AS}$, and then increases constantly until $T_B$. The same is seen in the ancestry shared among only EU samples, but where the initial increase was less rapid (due to a lower growth rate). Sharing between AS and EU is low, and only increases further back than $T_{EU-AS}$. On average, the relationship of both groups to AF genomes is indistinguishable, which is mirrored in the ancestry of AF genomes shared with genomes from either AS or EU. The ancestry shared among only AF genomes is higher (more recent) until reaching $T_B$, compared to the ancestry shared with either AS or EU, but then indistinguishable further back in time. Each comparison shows a gradient change at $T_{AF}$.

Ancestries inferred from true allele ages were overall consistent with patterns and times seen in the true ancestry profiles. For example, we see a rapid increase in the ancestry shared among only AS or only EU genomes until $T_{EU-AS}$, more recent shared ancestry among only AF genomes until $T_B$, and highly consistent gradients of the CCFs inferred between the different groups further back in time. However, we note that we infer artifacts at certain times, suggesting false changes in coalescent rates; for example at $\sim$2,500 generations ago among non-AF samples, but mostly in the distant past (>20,000 generations ago) and seen consistently across the whole sample. Such artifacts may result from incomplete information and variability of age-sorted sequences, as we only took a point estimate as the true age of mutations that may have occurred on relatively long branches.

For coalescent profiles inferred from estimated allele ages, we find notable differences between the values of the true and estimated fraction of the genome shared back in time. We also find artifacts that suggest false gradient changes, but again mostly limited to times in the distant past (>20,000 generations ago). The ancestry shared among genomes from the same population group increases rapidly in both AS and EU until $\sim$1,000 generations ago. Also, sharing between those groups is low until $\sim$1,000 generations ago, but which is slightly older than the actual time ($T_{EU-AS} = 848$ generations ago) of the split of AS and EU. Their split from AF is similarly shifted into the past, yet distinguishable from the ancestry of target genomes from AF shared with either AS or EU genomes. The ancestry of non-AF genomes shared with AF genomes is mirrored in the gradient and timing of the relationship between AF and non-AF genomes.

Overall, we note that the exact timings of events may not be reflected accurately by gradient changes along the inferred ancestry profiles, but we find the relative order of events to be consistent. Importantly, we find that ancestral relationships among and between different ancestry groups are qualitatively consistent.

#### S7 Estimation of variant age in publicly available data sets

We used GEVA to estimate the age of all variants on Chromosomes 1-22 (biallelic SNPs, except singletons and variants at alternate allele frequency >99%) in data from two human genome resources that are available in the public domain:

- 1000 Genomes Project (TGP), Phase 3, final release [3];
- Simons Genetic Diversity Project (SGDP), fully public data set [19].

The two data sets are described in detail further below. In total, we estimated the age of 45,393,705 variants across all autosomes, for each clock model (mutation clock, recombination clock, and joint clock). This includes 43,232,520 variants dated in sample data from TGP, and 15,834,824 variants from SGDP. For the 13,673,639 variants identified in both TGP and SGDP, we additionally estimated the age after combining information from both data sources (described in Section S7.4). Overall, we analyzed 32,087,462,147 haplotype pairs. A breakdown of variants dated per chromosome, as well as a summary of the haplotype pairs analyzed, is given in Table S1.

We make these results available as a public resource, referred to as the Atlas of Variant Age for the human genome. All data, including the full age estimation profiles for each clock model and the results of every pairwise analysis, are available online:

`https://human.genome.dating`

Throughout, we applied GEVA using the following specifications. We set $N_e$=10,000 to internally scale time in units of $2N_e$, which adheres to the usually quoted value [20]. Though, we note that more recent estimates of $N_e$ indicate a much lower effective size and high variability among different human populations [21, 22]. Results are reported in units of generations, after rescaling time given the specified value for $N_e$. We assumed a constant rate of mutation, set to $\mu = 1.2 \times 10^{-8}$ per base per generation, following recent estimates of the human mutation rate [23]. We used variable recombination rates according to HapMap genetic maps as available per chromosome (Phase 2; GRCh37) [2].

The maximum number of concordant and discordant haplotype pairs sampled per variant (specified by parameters $\max_{\mathcal{C}}$ and $\max_{\mathcal{D}}$) differed for the analyses conducted using TGP and SGDP data (see below). Throughout, variant age was estimated after applying the heuristic pair rejection method (Section S1.5) to exclude outliers in the pairwise TMRCA distributions of concordant and discordant pairs selected per variant, and we report the quality score ($QS$; Section S1.5.1) for the age estimated under each clock model.

##### S7.1 Information about ancestral and derived allelic states

The GEVA method (in its current implementation) assumes that ancestral and derived allelic states have been correctly assigned to the reference and alternate allele, respectively, as seen in a given data set. We acquired information as available for the human genome from Ensembl (human assembly GRCh37; release 92; version 20180221),* to determine the ancestral allele as predicted through multi-species alignments in the Ensembl *Enredo-Pecan-Ortheus* (EPO) pipeline.† We used this information to annotate available variant data, so as to (optionally) retain those variants in downstream analyses for which the ancestral allele was known and mapped to the reference allele. In TGP, we matched variant annotations based on chromosomal position, rsID, and consistent reference and alternate alleles. In SGDP, we did the same but omitted matching by rsID, as this information was absent in the available data set.

#### S7.2 Variants dated in 1000 Genomes Project (TGP) data

The final release TGP sample consists of 2,504 individuals (5,008 genomes) sampled from 26 populations worldwide.‡ We set $\max_{\mathcal{C}} = 500$ and $\max_{\mathcal{D}} = 500$ as the sampling limits for concordant and discordant pairs, respectively. For the 43.2 million variants dated across Chromosomes 1-22, we analyzed a total of 8.5 billion concordant and 21.3 billion discordant pairs, which involved the inference of the locally shared haplotype segment and the TMRCA posterior distribution at each pair. We recorded an overall computation time of approximately 2,699,936 hours (308 CPU years); measured as the sum of the system time elapsed per CPU core (Intel Xeon Gold 6126, Skylake SP, 2.60GHz).

We computed the quality score ($QS$) after rejecting outlier haplotype pairs for every variant and under each clock model; see Figure S23A. Estimation quality was high overall; for example, median $QS$ was $0.985$ for the joint clock model ($0.991$ mutation clock; $0.820$ recombination clock), and the proportion of variants with $QS > 0.95$ was 58.3% (joint clock; 61.0% mutation clock; 37.2% recombination clock). Because GEVA, by default, attempts to estimate the age of the alternate allele (assuming that it is derived and the reference allele is ancestral), we investigated differences in estimation quality arising from violations of this assumption, given the ancestral allelic states as predicted from the Ensembl EPO pipeline (Section S7.1); see Figure S23B. We found that 88.2% of variants had the ancestral allele as the reference allele; among those, median $QS$ was $0.993$ for the joint clock model ($0.996$ mutation clock; $0.846$ recombination clock). Among the 6.9% of variants where the ancestral allele matched the alternate allele, estimation quality was lower overall; median $QS$ was $0.645$ for the joint clock model ($0.680$ mutation clock; $0.568$ recombination clock). The relative proportion of variants with $QS > 0.95$ (joint clock) was 62.0%, 16.4%, and 52.2% for sites where the ancestral allele matched the reference, alternate, or neither allele, respectively. These results suggest that the quality score is informative as a measure of estimation quality, where low $QS$ values may indicate departures from baseline model assumptions in GEVA. However, we note that other sources of error exist, and that high $QS$ values may not validate estimation results.

* ftp://ftp.ensembl.org//pub/release-92/variation/vcf/homo_sapiens/homo_sapiens.vcf.gz
† https://www.ensembl.org/info/genome/compara/multiple_genome_alignments.html
‡ ftp://ftp.1000genomes.ebi.ac.uk//vol1/ftp/release/20130502/

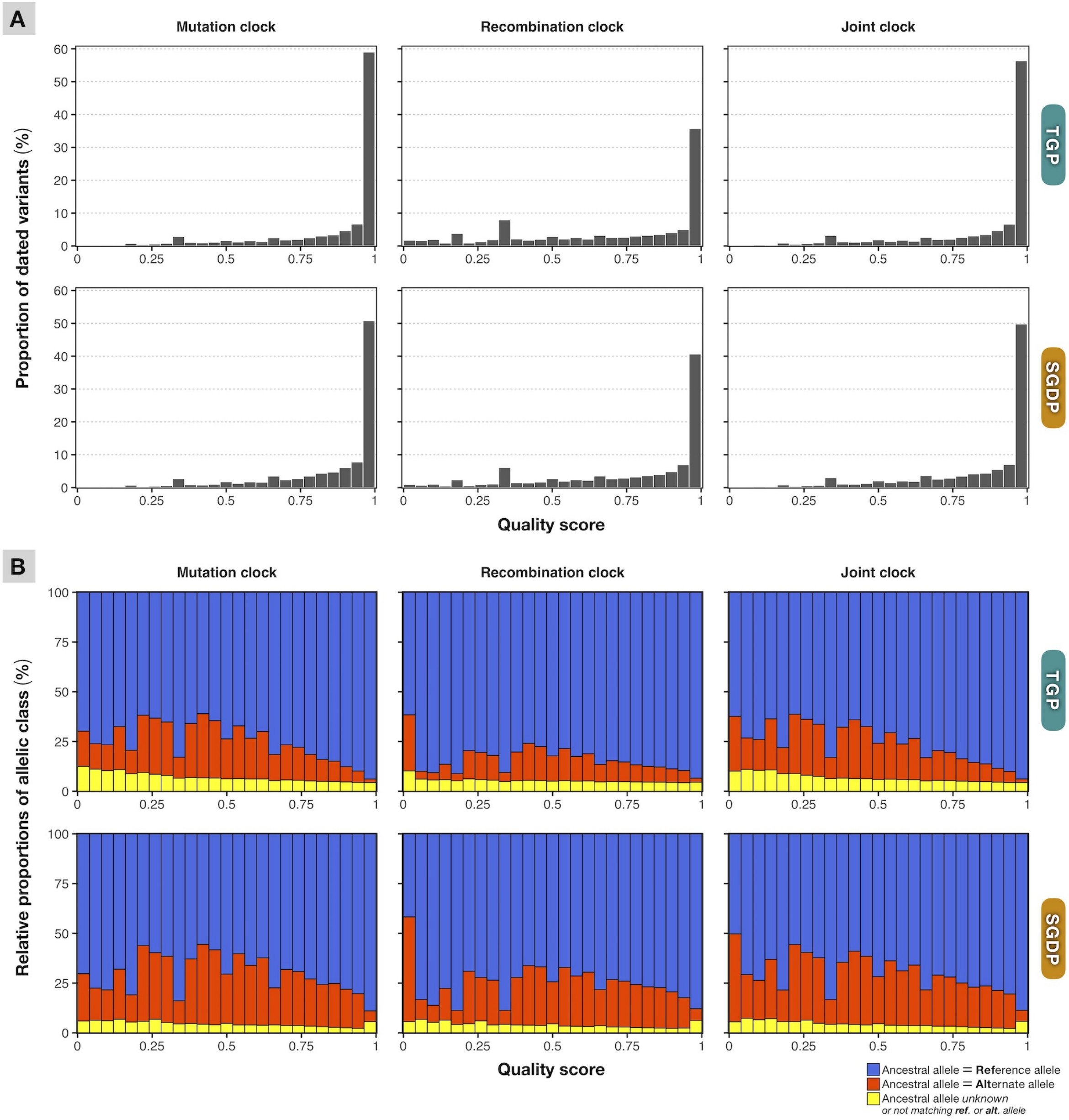

**Figure S23. Quality score for variants dated in TGP and SGDP data.** Panel **A** shows the histogram of the quality scores ($QS$) computed for variants dated in the 1000 Genomes Project (TGP; *top panels*) and the Simons Genome Diversity Project (SGDP; *bottom panels*), under each clock model. Histograms show the proportion of variants estimated with $QC$ values within 25 equally distributed bins, relative to the number dated in each data source; 43,232,520 variants in TGP and 15,834,824 in SGDP. Corresponding to (A), Panel **B** shows the relative proportions of allelic class found in each $QS$ bin; here defined as the ancestral allele matching the reference allele (*blue*), the ancestral allele matching the alternate allele (*red*), or where the ancestral allele was either unknown or did not match the reference or alternate alleles (*yellow*) at the variants (SNPs) dated in each data source. Note that GEVA attempts to estimate the age of the alternate allele by default, assuming that the reference allele represents the ancestral state.

###### S7.2.1 Pathogenic variants

We used information from the Ensembl Variant Effect Predictor (VEP; release 75) [6], available through TGP,* where functional consequences have been assigned to a subset of variants in the TGP sample (Phase 3; GRCh37). Variant effects have been predicted using PolyPhen-2 [7] and SIFT [8]. We selected all variants that had annotations from either PolyPhen-2 or SIFT and obtained their age from the Atlas of Variant Age. The set comprised 70,220 variants annotated by PolyPhen-2 and 67,539 variants annotated by SIFT, where 67,123 variants had annotations from both methods. The results shown in Figure S7 were generated on age estimates obtained from TGP data for the joint clock model, after excluding variants with low estimation quality ($QS \leq 0.5$) and where the reference allele did not match the ancestral allele.

##### S7.3 Variants dated in Simons Genome Diversity Project (SGDP) data

Data available from SGDP consists of 276 individuals (556 genomes) from 130 populations worldwide (fully public data set; hg19/GRCh37). We used the already phased panel (labelled "PS2") that had been phased using SHAPEIT2.† Given the relatively small sample size of the SGDP panel (compared to TGP), we set $\max_{\mathcal{C}} = 100$ and $\max_{\mathcal{D}} = 100$ as the sampling limits for concordant and discordant pairs per variant. We estimated the age for 15.8 million variants across autosomes, which involved the analysis of 0.7 billion concordant and 1.5 billion discordant pairs. Overall computation time was 33,168 hours (3.8 CPU years; Intel Xeon E5-2650 v2, Ivy Bridge EP, 2.60GHz). Note that the processing time was on average faster compared to the analysis of TGP data, due to the smaller sample size and the lower sampling limits set per variant.

We additionally re-estimated the age of >0.36 million variants on Chromosome 20 using $\max_{\mathcal{C}} = 500$ and $\max_{\mathcal{D}} = 500$, to assess differences resulting from estimation with lower and higher sampling limits. We found that age estimates were highly consistent under each clock model (Spearman's $\rho > 0.9$), indicating that the lower sampling limits ($\max_{\mathcal{C}} = 100$, $\max_{\mathcal{D}} = 100$) were sufficiently high for the main analysis of variants in SGDP; see Figure S24.

Estimation quality, measured after rejecting outlier haplotype pairs per variant and under each clock model, is shown in Figure S23A. For example, median quality score ($QS$) was $0.960$ for the joint clock model ($0.969$ mutation clock; $0.905$ recombination clock), and the proportion of variants with $QS > 0.95$ was 51.8% (joint clock; 53.1% mutation clock; 42.8% recombination clock). Differences in quality with regards to the ancestral state of the reference or alternate allele are shown in Figure S23B. As in the analysis of variants in TGP, the majority of variants in SGDP (81.5%) had the ancestral allele as the reference allele, for which median $QS$ was $0.980$ (joint clock; $0.980$ mutation clock; $0.929$ recombination clock). However, compared to estimation in TGP data, we found that a higher proportion of variants had the ancestral allele as the alternate allele (14.0%), and differences in quality were not

* ftp://ftp.1000genomes.ebi.ac.uk//vol1/ftp/release/20130502/supporting/functional_annotation/
† https://sharehost.hms.harvard.edu/genetics/reich_lab/sgdp/phased_data/PS2_multisample_public/

as pronounced as in TGP; median $QS$ was 0.794 (joint clock; 0.816 mutation clock; 0.737 recombination clock). The relative proportion of variants with $QS > 0.95$ (joint clock) was 56.2%, 22.3%, and 64.1% for sites where the ancestral allele matched the reference, alternate, or neither allele, respectively. Differences in estimation quality measured for variants dated in TGP and SGDP may also arise from the different sampling limits applied. Note that we used external data to determine the ancestral allele (Ensembl EPO pipeline; Section S7.1), and that we included this information only at sites where genomic position and bases at both alleles matched unambiguously. For variants identified in both TGP and SGDP, we found that age estimates independently obtained from the two data sources agreed (see results in Figure S5).

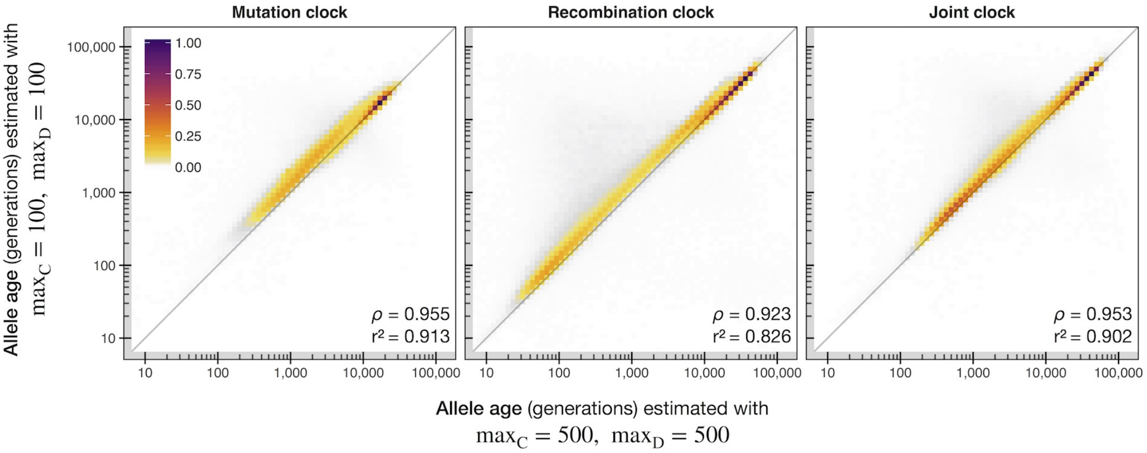

**Figure S24. Consistency of variant age estimated with high and low sampling limits.** Density scatterplot showing the relationship between allele age estimated with different sampling limits for concordant and discordant pairs; set to $\max_{\mathcal{C}} = 500$ and $\max_{\mathcal{D}} = 500$ (*x-axis*), and set to $\max_{\mathcal{C}} = 100$ and $\max_{\mathcal{D}} = 100$ (*y-axis*). Age was estimated for all variants on Chromosome 20 in SGDP (>0.36 million) under each clock model. Colors indicate the relative density scaled by the maximum per panel. The inserts (*bottom*) show the correlation between the two analyses; Spearman rank correlation coefficient ($\rho$) and the square of the Pearson correlation coefficient ($r^2$; calculated on log-scaled allele ages).

#### S7.4 Combined age estimation of variants present in TGP and SGDP

The true age of a variant refers to the time of a singular mutation event in the past, which follows from the assumption (infinite sites model) that mutations occur only once per genomic locus in the history of the population. Although this assumption is readily violated in reality, in particular for very old mutations or if the mutation rate is high, we would nonetheless expect it to hold for the vast majority of variants observed in the (human) genome. Estimating the age of the same allele in different data sources, in which it has been identified in genomic sequence data of independent (unrelated) samples, should therefore yield consistent results.

We identified 13,673,639 variants present as biallelic SNPs (non-singletons and below 100% allele frequency) in both TGP and SGDP; matched by genomic position and reference and alternate alleles. The ages of alleles at corresponding sites, independently estimated from TGP and SGDP data, were highly consistent; see Figure S5. We then, additionally, estimated the age of these variants by combining information from both sources, to implicitly increase the genealogical resolution, but without combining sequence data directly.

Recall that we estimate the age from the composite distribution of locally inferred TMRCA posteriors at concordant and discordant haplotype pairs; see Section S1.4, Equation (27). The TMRCA posterior is modeled using the Gamma distribution, where parameters $\alpha, \beta$ are obtained from the data as defined by the clock model used (mutation clock, recombination clock, or joint clock). For each variant, we combined information by recovering TMRCA posteriors from the parameter values as obtained in the two data sources (without rejecting outliers). We estimated the "combined" age, and computed a quality score, after rejecting outlier pairs in the combined sets of concordant and discordant comparisons.

### S8 Inference of the ancestry shared between individuals and populations

We used the dynamic programming method presented in Section S2 to infer the cumulative coalescent function (CCF) between every pair of genomes within TGP, as well as SGDP. Inference of the ancestry shared between individual genomes was conducted separately on data from Chromosomes 1-22. Information from the Atlas of Variant Age was used to generate, in each analysis, the age-sorted sequence of observed allele sharing between the haploid chromosomes of a given target and comparator individual. Throughout, we used age estimates generated from the joint clock model, for variants dated in TGP, as well as SGDP. We considered only those variants where the ancestral allele matched the reference allele, and where estimation quality was reasonably high ($QS > 0.5$). The fraction of the genome shared was inferred at a resolution of $S = 200$ states (see Section S2.1). We approximated the CCF over a fixed grid of 1,000 time points between 10 generations and 1 million generations ago (evenly distributed on log-scale). A summary profile was generated per diploid individual by aggregating individual CCFs as the weighted average across chromosomes, using the number of variants retained per chromosome to calculate weights (as described in Section S2.4).

The coalescent profiles inferred between each chromosomal pair in TGP and SGDP, as well as reduced summaries of the ancestry shared with the whole sample per individual, are publicly available online at https://human.genome.dating/ancestry/index.

**Ancestry sharing in TGP.** Of the 43.2 million variants dated in TGP, we retained 34.4 million across autosomes after filtering. We inferred the CCF between the haploid genomes from the 2,504 individuals available in the TGP sample, as well as the panel of 31 related individuals that were excluded from the final release data set.* Individuals in this additional panel comprised trios, parents, siblings, and second order relatives, who were part of the initial data set and also used to produce the variant call set of the final data set. Thus, we inferred the CCF for all 5,070 haploid chromosomes as target genomes, in turn against all others as comparator genomes, resulting in 25.7 million pairwise coalescent profiles per chromosome, and 6.3 million summary profiles after aggregating individual CCFs.

**Ancestry sharing in SGDP.** Coalescent profiles were inferred on 11.7 million variants across autosomes, retained after filtering from the full set of 15.8 million variants dated in SGDP. We inferred the CCF between each pair of haploid chromosomes in the sample of 278 diploid individuals, resulting in 308,580 pairwise coalescent profiles per chromosome, and 77,284 summary profiles after aggregating individual CCFs. Additionally, we prepared cross-chromosome summaries for groups of individuals, by aggregating CCFs among individuals belonging to the 130 ancestry groups. This resulted in 16,900 summary profiles that describe the ancestry shared between each demographic unit.

* ftp://ftp.1000genomes.ebi.ac.uk//vol1/ftp/release/20130502/supporting/related_samples_vcf
